# Return of the GEDAI: Unsupervised EEG Denoising based on Leadfield Filtering

**DOI:** 10.1101/2025.10.04.680449

**Authors:** Tomas Ros, Victor Férat, Yingqi Huang, Cristina Colangelo, Seyed Mostafa Kia, Thomas Wolfers, Serge Vulliemoz, Abele Michela

## Abstract

Current electroencephalogram (EEG) denoising methods struggle to remove the complex physiological and environmental artifacts typical of real-world settings, which both hinders the isolation of true neural activity and limits the technology’s translational potential. We present the Generalized Eigenvalue De-Artifacting Instrument (GEDAI), a novel algorithm for denoising highly contaminated EEG. GEDAI employs *leadfield filtering* to selectively remove noise and artifacts that diverge from a theoretically defined EEG forward model. This approach offers unique advantages over existing solutions, including 1) denoising of highly corrupt recordings without “clean” reference data, 2) single-step correction of artifactual epochs and bad channels, 3) unsupervised detection of brain and noise components based on the signal and noise subspace alignment index (SENSAI). In ground-truth simulations with synthetic and empirical EEG contaminated with realistic artifacts (EOG, EMG, noise), GEDAI globally outperformed leading denoising techniques based on principal component analysis (ASR) and independent component analysis (IClabel, MARA), revealing large effect sizes in challenging scenarios with simultaneous artifact mixtures, low signal-to-noise ratio (-9 dB), and high temporal contamination (up to 100%). Its superior denoising also enhanced neurobehavioral predictions, yielding highest accuracies in ERP classification and brain fingerprinting. GEDAI’s autonomy, computational speed and noise-resilience could find future applications in 1) real-world *medical*, *mobile* and *dry* electrode EEG recordings 2) magnetoenecephalography (MEG) denoising (given the shared M/EEG forward model), and 3) *real-time* brain-computer interfaces (BCIs). The Matlab code for GEDAI is available as an open-source EEGLAB plugin at https://github.com/neurotuning/GEDAI-master

## Introduction

Despite being well-researched and affordable, the multi-channel electroencephalogram (EEG) struggles to expand beyond controlled research and clinical settings (Sawangjai et al., 2020). A major hurdle to broader EEG use is contamination by non-cerebral signals, known as artifacts (Mumtaz et al., 2021). EEG artifact removal often requires the supervision of a trained operator, leading to reduced scalability and reliability. Hence, tackling artifacts has recently been rated by experts as the most important development needed in EEG research (Mushtaq et al., 2024).

In the realm of medical applications (Fratangelo et al., 2025), EEG artifacts can obscure or mimic genuine brain activity, leading to inaccurate diagnoses and compromised patient care (e.g. in epilepsy, intensive care or sleep monitoring) (Amin et al., 2023). Furthermore, artifacts significantly hinder the development and reliability of brain-computer interfaces (BCIs) (Mak & Wolpaw, 2009), which rely on consistent and accurate decoding of neural signals to translate brain activity into commands for external devices or neurofeedback (Ros et al., 2014). Artifacts introduce noise that can disrupt this decoding process, leading to reduced accuracy, slower response times, and ultimately, a less effective BCI system. Therefore, there remains a critical need for a fast, fully automated, and noise-agnostic EEG method to reliably remove all types of artifacts, particularly in highly contaminated recordings where clean reference data is unavailable.

Several previous strategies have been developed to clean multi-channel EEG signals. The most popular of which is independent component analysis (ICA), which performs blind source separation and can be used to remove temporally-independent components from the signal. While effective at removing ocular (Jung et al., 2000) and muscular artifacts (Olbrich et al., 2011), ICA is computationally demanding, requires a case-by-case inspection of components to reject that calls for expert human input or that from machine learning classifiers (Frølich et al., 2015; Radüntz et al., 2017). Moreover, ICA cannot inherently identify Gaussian distributed noise, which might be spread across the components or remain as unexplained residual variance . ICA is therefore not an ideal choice for inexperienced users, large datasets or online analysis (despite promising attempts e.g. Hsu et al., 2014).

Another algorithm, called artifact subspace reconstruction (ASR), directly uses principal component analysis (PCA) to exclude EEG components exceeding a variance threshold, but requires a portion of clean EEG data as a reference (Kothe & Jung, 2015). While faster, relying solely on a variance threshold is not per se sufficient to distinguish between signal and noise (de Cheveigné & Parra, 2014), as in the case when noise amplitude is below that of the signal. Moreover, since ASR is based on PCA, it can only separate orthogonal components (Cohen, 2022a), which may limit its effectiveness when trying to resolve complex artifact mixtures.

To overcome the limitations of ICA and PCA, Generalized Eigenvalue Decomposition (GEVD) offers a promising alternative (Koles, 1991; Y. Wang et al., 1999). GEVD works by taking a pair of EEG covariance matrices and decomposing them jointly – a process known as "joint diagonalization" – to find common underlying components (de Cheveigné & Parra, 2014). The core objective of this joint diagonalization is to find a set of components that maximizes the variance ratio between the signal and reference matrices (Cohen, 2022a). Another key advantage is that the resulting GEVD components do not need to be orthogonal (unlike PCA) or conform to Gaussian assumptions (unlike ICA) (Cohen, 2022a; Parra & Sajda, 2003).

GEVD is widely employed in the BCI field for mental classification tasks, where it is known as the common spatial patterns (CSP) algorithm (Blankertz et al., 2008; Lotte & Guan, 2011). It has also been proposed for brain source separation (Cohen, 2022a; Parra & Sajda, 2003), correction of ocular artifacts (Gouy-Pailler et al., 2009), removal of stimulation artifacts (Haslacher et al., 2021) and generic M/EEG data cleaning (Boudet et al., 2012; Somers et al., 2018; F. Wang et al., 2025).

We refer to this work as the “return” of the GEDAI (Generalized Eigenvalue De-Artifacting Instrument) because, while GEVD has been used in EEG denoising before, our approach revisits it by addressing key limitations. An existing challenge in applying GEVD is the need to specify the reference covariance matrix (refCOV) for decomposition. This typically involves selecting empirical EEG segments deemed "clean" (Haslacher et al., 2021; F. Wang et al., 2025) or "artifactual" (Gouy-Pailler et al., 2009; Somers et al., 2018). However, this process is often subjective and circular since identifying representative segments requires prior knowledge of their typical characteristics. GEDAI addresses this by constructing the refCOV theoretically, deriving it from EEG signal generation principles using the forward model via the leadfield matrix (Weinstein et al., 2000). This leadfield-based refCOV represents the expected EEG signature of brain activity generated by internal sources, offering a principled way to define the "clean signal" subspace for decomposition. We refer to this forward-modeling approach to separate brain signals from noise as leadfield filtering (LFF). An additional challenge with GEVD is determining the cutoff that separates ’signal’ (brain activity) from ’noise’ (artifacts) components. GEDAI addresses this by using the theoretical refCOV as a benchmark, selecting the optimal threshold based on subspace similarity between the cleaned EEG covariance and the refCOV (see **Methods** section).

To evaluate GEDAI’s performance, we tested it against prominent artifact correction techniques using both synthetic and empirical EEG data, following recommendations for robust testing (Mumtaz et al., 2021). This comparison includes the fast PCA-based method ASR (Kothe & Jung, 2015) and two widely used ICA-based methods known for effective results: ICLabel (Pion-Tonachini et al., 2019) and Multiple Artifact Rejection Algorithm (MARA) (Winkler et al., 2014). This comparative analysis aims to position GEDAI relative to current state-of-the-art methods.

## Methods

**Section I** of the methods describes the mathematical details of the GEDAI denoising algorithm, whose code we also release as an open plugin for the EEGlab toolbox in MATLAB (Delorme & Makeig, 2004).

**Section II** of the methods describes benchmarking using ground-truth simulations, where GEDAI’s performance is compared to current state-of-the-art algorithms for EEG de-artifacting, including ASR (Kothe & Jung, 2015), ICLabel (Pion-Tonachini et al., 2019) and MARA (Winkler et al., 2014). The simulations were performed using both synthetic and empirical EEG datasets to which noise and/or artifacts were added.

**Section III** of the methods describes each algorithm’s denoising performance in the context of neurobehavioral prediction. Here, two publicly-available EEG datasets were used to compare how the denoising algorithms influence the prediction accuracy of sensory stimuli (“visual oddball”) and individual identity (“brain fingerprinting”).

### Section I: The GEDAI algorithm

As illustrated in **Figure 1A** above, multi-channel EEG may be considered to be a linear summation of electrical activities from a brain “signal” subspace with one containing different types of non-cerebral noise or “artifact”. This mixture may be “unmixed” by linear decomposition techniques (e.g. PCA or ICA) into separate components with individual source locations (“topographies”) and respective time-courses (“waveforms”). However, as "blind" source separation methods, PCA and ICA leverage statistical properties within mixed data to recover underlying sources, functioning without *a priori* knowledge of the original signals or their mixing process. GEDAI combines prior knowledge of the brain’s “signal” subspace (i.e. its spatial covariance) with generalized eigenvalue decomposition (GEVD) to more effectively separate source components belonging to the artifact-subspace from those of the brain- subspace.

**Figure 1.**
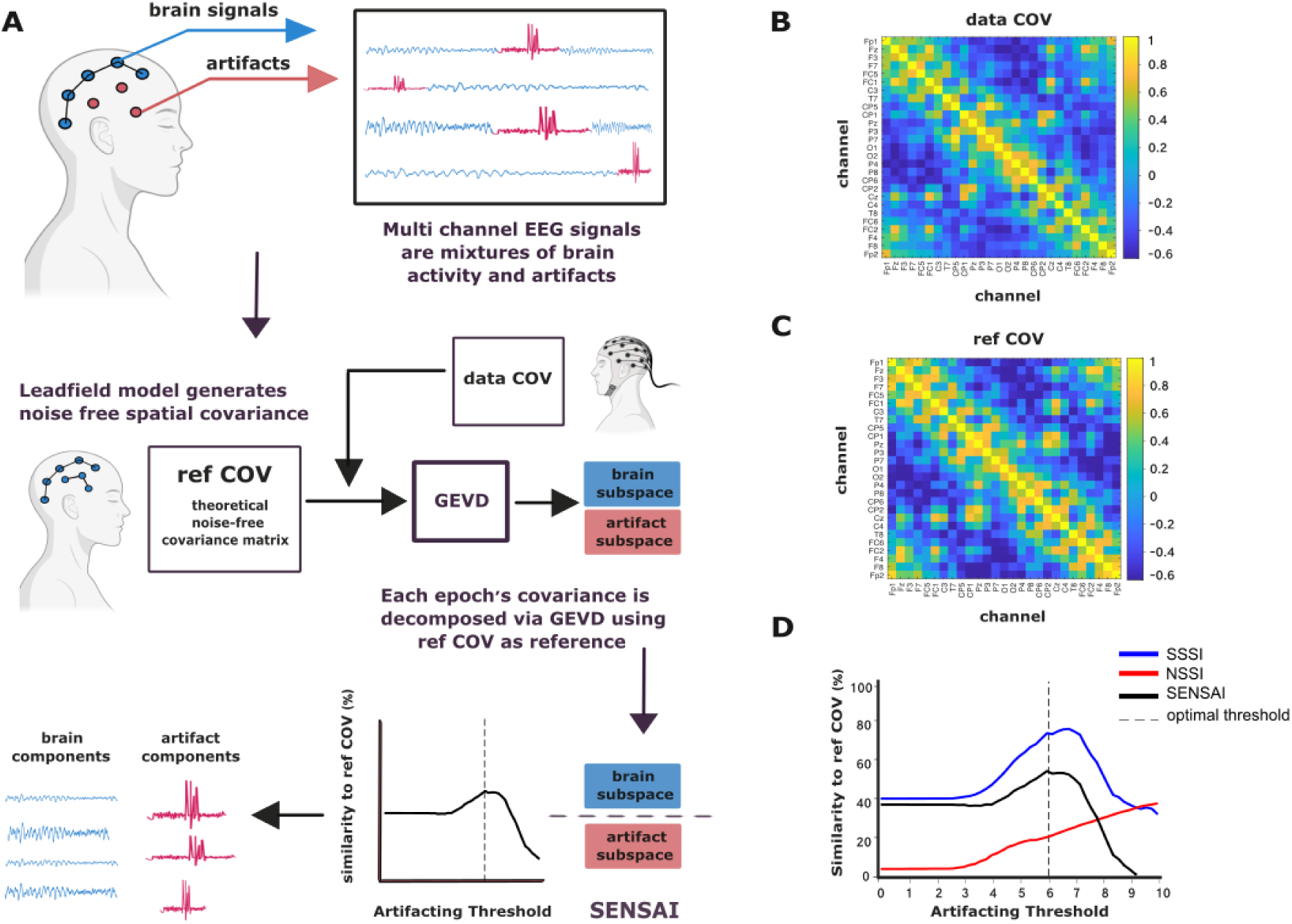
An overview of the GEDAI algorithm A) Schematics illustrating the GEDAI pipeline; dataCOV: empirical data covariance matrix; refCOV: theoretical covariance matrix from a leadfield model. B) An example dataCOV of clean EEG data. C) An example refCOV of a leadfield matrix. D) Graph illustrating similarity between the denoised data and the refCOV across a range of artifact thresholds. SSSI: Signal Subspace Similarity Index, blue curve; NSSI: Noise Subspace Similarity Index, red curve; SENSAI: SSSI - NSSI, black curve; vertical dashed line: optimal artifacting threshold.

First, refCOV is derived from a precomputed leadfield matrix for standard 10-5 system electrode locations or, for non-standard electrode locations, from an interpolated leadfield matrix calculated ‘on-the-fly’ via spherical interpolation. Next, the whole input EEG signal (including artifacts) is epoched in circa 1-second windows, and an individual data covariance matrix (dataCOV) is generated for each epoch. Then, each data covariance matrix is decomposed with GEVD, using refCOV as a fixed reference matrix across all epochs. After, the output EEG data is reconstructed and evaluated using the Signal & Noise Subspace Alignment Index (SENSAI), in order to determine the optimal cutoff that separates brain from artifactual components, sweeping across a range of artifacting strengths. By respectively maximising and minimizing the subspace similarities of the retained and removed data with the refCOV, across different thresholding strengths, GEDAI estimates the optimal cutoff for component removal. For the final step, the denoised time-series of each epoch is then reconstructed by using only the GEVD components belonging to the brain-subspace.

#### Estimation of the refCOV

The GEDAI EEGlab plugin offers two options for refCOV estimation. The first method uses a pre-computed covariance matrix of 343 standard EEG electrode locations (10-5 system), from which the plugin automatically matches the electrode labels present in the EEG recording (e.g. Fp1, Pz, etc.). This leadfield matrix was generated with the Brainstorm Toolbox (Tadel et al., 2011) using the OpenMEEG algorithm (Gramfort et al., 2010) and the ‘fsaverage’ adult head model (FreeSurfer’s default template based on 40 normative brains), employing the Boundary Element Method (BEM) with 3630 unconstrained brain dipolar sources (1210 vertices × 3 orientations).

For non-standard EEG recording montages, there is a second option: spherically interpolating the precomputed leadfield to custom electrode locations. This method requires that the electrodes’ spatial coordinates are provided within EEGlab. Although slightly less accurate, this approach is computationally much faster than estimating a custom BEM leadfield, taking only a couple of seconds compared to the several minutes required to recompute with OpenMEEG.

The leadfield matrix parametrizes the “forward model” of how the EEG is generated by sources of neuronal activity in the brain, i.e., the EEG_activity with dimensions [channels x time], the leadfield_matrix with dimensions [channels x sources], and the brain_source_activities with dimensions [sources x time]:

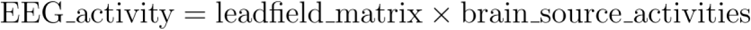

From this, the spatial covariance matrix, refCOV, with dimensions [channels x channels] can be simply computed as:

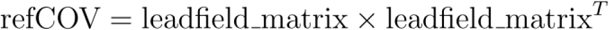

GEDAI’s source-space to electrode-space projection relies on two main theoretical assumptions (de Munck et al., 1988, 1992): electrode potential is a weighted sum of underlying dipolar sources, and each source’s strength is independent of other source parameters. Under these conditions, de Munck et al. demonstrated a linear relationship between electrode signal covariance and the summed, weighted covariance of individual dipolar sources.

We provide a simple empirical example supporting the aforementioned assumptions in **Figure 1** where a leadfield-generated covariance matrix (panel C) may be qualitatively compared to the covariance matrix calculated from clean EEG data (panel B) (Subject #2 from the Ehrlich dataset, see Methods). Both matrices were normalised [channel x channel] (i.e. zero mean with a standard deviation of one) for visualisation.

#### Generalized Eigenvalue Decomposition

Threshold-based artifact rejection methods rely on the principle that artifacts are usually large and relatively rare events occurring in the signal. Artifacts can therefore be identified as appearing in the upper tail of the eigenvalue distribution of EEG signals, and be excluded based on a defined cutoff (Lazarevic & Kumar, 2005; Li et al., 2023). The large magnitude of artifacts is exploited by PCA-based procedures such as ASR, but these are constrained by orthogonality and blind to the spatial origin of the signal components.

In contrast, GEDAI employs GEVD to isolate and remove artifactual components from the signal. However, GEVD alters the significance of the largest eigenvalues. Instead of representing "components with high variance," they now signify "components with high variance that *maximally* deviate from the reference covariance matrix (*refCOV*)." Given that the *refCOV* encodes components that spatially originate only within the brain, this *a priori* provides GEVD with extra “supervision” for separating artifactual from neural sources.

The GEVD may be summarised in one line of MATLAB/Python code, where *e* is the epoch number:

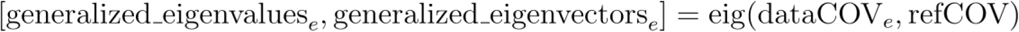

Mathematically, the GEVD decomposition follows the linear algebra equation:

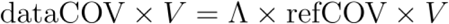

Where dataCOV is the covariance matrix of a single epoch extracted from the EEG data, *refCOV* is the predetermined reference covariance matrix, Λ is the diagonal matrix containing the generalized eigenvalues and V is the matrix containing the generalized eigenvectors. In GEDAI, a regularization technique is applied to *refCOV* before GEVD to enhance numerical stability, manage singular matrices, and reduce overfitting (Cohen, 2022a).

#### The SENSAI algorithm: separating neural from artifactual components

Akin to finding the boundary that separates oil from water, differentiating artifacts from neural signals depends on the exact magnitude of the eigenvalues, represented by the threshold *T.* SENSAI (Signal & Noise Subspace Alignment Index) essentially estimates the correct threshold *T* by evaluating the “improvement” in the output EEG signal quality by sweeping over multiple threshold values of *T*. Here, the output EEG signal quality is represented by the SENSAI score, which is proportional to the cosine similarity between the *refCOV* matrix and the empirical covariance of the retained (i.e. *Signal*) and the removed (i.e. *Noise*) data.

As can be seen in **Figure 1D**, the SENSAI function computes a similarity index of the retained and removed EEG data for each threshold tested. The similarity index is calculated first by taking the covariance matrix of the cleaned EEG data and the *refCOV*, and performing a classical eigendecomposition separately on both (i.e. equivalent to PCA). The principal angles between the top components (i.e. eigenvectors) of the *refCOV* and cleaned data subspaces are then estimated (Knyazev & Argentati, 2002). Here, given two subspaces with their orthonormal bases *A* (from *refCOV*) and *B* (from cleaned data), the principal angles θi between these subspaces can be computed as follows:

Compute the projection matrix C:

Perform Singular Value Decomposition on the projection matrix :

where *W* is related to the orthonormal basis of the subspace *A*, *Q* is related to the orthonormal basis of the subspace *B*, and Σ is a diagonal matrix containing the cosines of the principal angles between the two subspaces. The principal angles θi are obtained from the singular values σi in the diagonal matrix Σ:

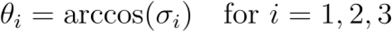

To estimate the Signal Subspace Similarity Index (SSSI), we take the cosine of each principal angle and multiply them together:

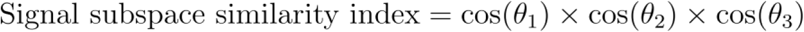

Where 𝜃1, 𝜃2 and 𝜃3 are the principal angles between the top 3 eigenvectors of the cleaned data and the top 3 eigenvectors of the refCOV. Similarly, the Noise Subspace Similarity Index (NSSI) is also estimated between the top 3 eigenvectors of the residual noise data (removed from the EEG) and the refCOV. By estimating the SSSI and NSSI through a broad range of artifacting thresholds, it becomes possible to optimize the tradeoff between either removing or retaining too many artifactual components. For example, increasing the threshold strength at first increases the similarity of the cleaned EEG covariance with *refCOV*, as non-cerebral components are initially removed. Past a certain point, however, the threshold becomes too aggressive and cerebral activity starts to be removed from the EEG recording, thus reducing the similarity of the cleaned EEG covariance with refCOV. Similarly, by calculating the NSSI for the data being removed from the EEG in the cleaning process, we can quantify how much potentially cerebral EEG activity gets included as noise.

SENSAI calculates a tradeoff score, called the SENSAI score, by subtracting the subspace similarity index of the noise from the similarity index of the cleaned EEG, or mathematically:

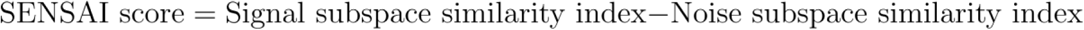

The optimal threshold for cleaning the data is simply the eigenvalue that corresponds to the *maximum of* this score.

#### Denoised signal reconstruction

Cleaned EEG signal reconstruction occurs epoch-by-epoch. Within each epoch, artifactual components are first identified (eigenvalues > T), and a spatial filter targeting these artifacts is formed using the corresponding eigenvectors. Concurrently, the activation patterns (or inverse spatial filters) for these artifactual components are derived from the pseudo-inverse of the eigenvector matrix. The artifactual activity is then estimated in sensor space by combining these activation patterns with their respective time courses (calculated using the spatial filter and the original EEG). Finally, subtracting this reconstructed artifact activity from the original epoch yields the denoised EEG signal. To prevent signal discontinuities that can arise from concatenating independently processed EEG epochs, GEDAI employs a 50% overlapping window approach during its epoch-by-epoch reconstruction. These overlapping segments are combined using cosine weighting to ensure smooth transitions at epoch boundaries, mitigating the edge effects inherent in segmented processing.

#### Multiresolution GEDAI

Although GEDAI can be applied end-to-end solely to broadband data (e.g. 1-45 Hz), we have found that multiresolution-based denoising performs significantly better on pilot data. Multiresolution analysis decomposes the signal into frequency sub-bands across multiple scales, which permits perfect reconstruction, ensuring lossless recovery of the original EEG data from its components (Clark et al., 1995). Hence, following an initial GEDAI denoising step on broadband EEG to remove the largest artifacts, the GEDAI cleaning algorithm is subsequently applied to spectrally decomposed EEG data using a maximum overlap discrete wavelet transform (MODWT). The benefit of this extra step comes from the fact that artifact distributions clearly differ between EEG bands (ranging from delta to gamma). Thus, frequency-specific thresholds are needed to ensure that high-frequency artifacts (e.g. in the gamma band) are not rejected using the same eigenvalue threshold estimated for lower frequency bands (e.g. the delta band). Hence, the GEDAI algorithm consists of a secondary step on MODWT decomposed signals filtered in 10 dyadic frequency bands, using the ‘Haar’ wavelet. Among the wavelet families, the Haar wavelet has the best temporal localization, which is an important property for pinpointing abrupt changes, discontinuities, or spikes typical of artifactual signals. Compared to other types of wavelet analysis, the MODWT offers advantages like conservation of signal energy (perfect reconstruction) and time alignment (zero-phase filtering, shift invariance) across decomposition levels. For an EEG recording sampled at 256 Hz, the wavelet bands produced by the MODWT are as follows: 64-128 Hz (band 1), 32-64 Hz (band 2), 16-32 Hz (band 3), 8-16 Hz (band 4), 4-8 Hz (band 5), 2-4 Hz (band 6), 1-2 Hz (band 7), 0.5-1 Hz (band 8), 0.25-0.5 Hz (band 9), 0.125-0.25 Hz (band 10). Then, for each narrow-band the optimal artifacting threshold is estimated separately via the SENSAI algorithm, and the final clean signal is reconstructed by adding up all the denoised wavelet bands.

### Section II: Benchmark EEG simulations

Using EEGlab v2025.0.0 we compared the GEDAI algorithm to 3 state-of-the-art (SOTA) denoising algorithms (ASR, MARA and ICLabel) in terms of ground-truth signal reconstruction on both synthetic and empirical EEG data.

#### Denoising algorithms: ASR, IClabel, MARA, GEDAI

Three established automated denoising algorithms were used to benchmark GEDAI, namely ASR, IClabel and MARA. Like GEDAI, these are all projection algorithms that “correct” artifacts by subtracting them from EEG data, and which technically incur no temporal data loss. A panoply of pipelines also exists for denoising EEG data that involve wholesale “rejection” of EEG segments, but these are designed to solve a different problem, where full recovery of the original EEG signal is not the objective (Bailey et al., 2023; Gabard-Durnam et al., 2018; Hajhassani et al., 2024). We have also not included deep learning based denoising algorithms (X. Zhang, 2024), as these techniques rely on extensive training datasets and can exhibit limited generalizability when applied to out-of-sample data. This could be investigated in future work and is beyond the scope of this paper.

#### Raw

As a sanity-check, we provide the “raw” unprocessed data for comparison, which was used as the direct input to each denoising algorithm. Hence, the *raw* condition did not contain any denoising and/or preprocessing, and consisted of the original EEG contaminated with artifacts.

#### ASR

As the first denoising algorithm, we included ASR (i.e. EEGlab’s *clean_ra*w_data v2.91), as it is a fast, real-time capable method for EEG cleaning (Kothe, 2013). The ASR algorithm performs a standard eigendecomposition using one covariance matrix as input (i.e. PCA), instead of a generalized eigendecomposition that utilizes two covariance matrices as input (i.e. GEVD). We used EEGlab’s ASR plugin default burst criterion value of 20 SD (Chang et al., 2020). Given that bad (i.e. noisy) channels negatively affect ASR performance, we activated *clean_raw_data*’s bad channel detection only for artifact categories where noisy channels were present, i.e. NOISE or NOISE + Electromyography (EMG) + Electrooculography (EOG). Since bad channel detection was not universally applied over all artifact categories, the benchmark was not fully "noise-agnostic” and slightly advantaged ASR.

#### IClabel

The second denoising algorithm was ICLabel v1.6 (Pion-Tonachini et al., 2019). This ICA-based technique can automate artifact rejection by evaluating ICA topographies with a machine learning based classifier trained on ICA topographies from over 6,000 datasets. Here, the Infomax-extended ICA algorithm (Lee et al., 1999) was used based on its superior performance compared to other ICA algorithms (Delorme et al., 2012). Firstly, given that bad channels may impair the quality of ICA decomposition, the ICA was preceded by EEGlab’s *clean_raw_data* bad channel rejection step only for artifact categories where noisy channels were present (i.e. NOISE or NOISE + EMG + EOG). Moreover, if the artifact category contained only EMG or EOG artifacts, IClabel was configured to remove only those ICA components identified as EMG or EOG artifacts (all other components were retained). On the other hand, if the artifact category contained noise (i.e. NOISE or NOISE + EMG + EOG), IClabel was configured to only retain components that were classified as ‘brain’ or ‘other’. Since the above settings were not universally applied over all artifact categories, the benchmark was not fully "noise-agnostic” and provided a marginal advantage to IClabel.

#### MARA

For the third denoising algorithm, we selected MARA v1.2, the Multiple Artifact Rejection Algorithm (Winkler et al., 2014). This ICA-based technique also involves a machine learning classifier trained on a large database of expert-labeled artifactual topographies. MARA is integrated within popular denoising pipelines, such as the Harvard Automated Processing Pipeline for Electroencephalography (HAPPE) (Gabard-Durnam et al., 2018). Unlike IClabel, MARA directly classifies ICA components as artifactual (or not), and hence only those components were removed. To facilitate comparisons to IClabel, the Infomax-extended ICA algorithm was used (Lee et al., 1999), which was preceded by EEGlab’s *clean_raw_data* bad channel rejection step only for artifact categories where noisy channels were present (i.e. NOISE or NOISE + EMG + EOG). Since bad channel detection was not universally applied over all artifact categories, the benchmark was not fully "noise-agnostic”, which slightly favoured MARA’s performance.

#### GEDAI

We executed EEGlab’s GEDAI v1.0 plugin using its default settings: with denoising strength set to ‘auto’, an epoch size of 1.0 seconds, and a ‘precomputed’ BEM lead field matrix. No bad channel rejection step was applied to any of the artifact categories. The above settings were universally applied across all datasets, hence the GEDAI benchmark results can be considered as effectively "noise-agnostic".

#### Denoising scenarios: signal-to-noise ratio, temporal contamination, artifact type

Each denoising algorithm was run with its *default* parameter settings across the complete range of noise scenarios described below. Fixing the parameter settings was necessary to test each algorithm’s automation and/or generalization to a range of real-world “noise” scenarios, which were varied across 3 key axes:

- ● ***Signal to Noise Ratio (SNRbefore)*** of clean EEG power relative to artifact power (-9, - 6, -3, 0 dB)

where SNRbefore = 10 * log10(original_clean_signal_power / original_artifact_power) was estimated across all epochs (i.e. the whole EEG recording)

- ● ***Temporal contamination*** of EEG contaminated by artifacts (25, 50, 75,100%) where % reflects the proportion of samples containing artifacts. For example, for a

dataset of 60 seconds and sampled at 200 Hz, we added artifact segments with random offsets into a total of 4000, 6000, 8000, or all 12000 samples. The artifact segment duration was randomized, ranging from 1 sample to 1 second..

- ● ***Artifact type*** (EOG, EMG, NOISE, or their combination), where different artifact types were linearly superimposed within each epoch.

### Denoising Benchmark: Synthetic and Empirical EEG Simulations

#### Synthetic EEG simulations

Synthetic data for these simulations were generated using a combination of the EEG forward modelling toolbox SEREEGA (Krol et al., 2018) and the *EEGdenoiseNet* dataset based on real EOG and EMG electrode recordings (H. Zhang et al., 2021).

#### Clean EEG signal generation

Here, we exclusively used the SEREEGA toolbox to generate clean “background” EEG data. A total of 10 independent datasets were simulated with distinct spatio-temporal dynamics, including: 2 datasets with autoregressive noise, 6 datasets with varying combinations of colored noise (white, blue, brown) plus amplitude-modulated oscillations (in delta, theta, alpha and beta bands), and 2 datasets with simulated epileptic spiking activity (small or large spikes). The forward model consisted of the New York head model (i.e. ICBM152 average template) with a precomputed Finite Element Method (FEM) leadfield (Huang et al., 2016). Here, 100- 300 dipolar sources, randomly distributed within the brain volume, were used to generate a multi-channel time series across 100 independent 1-second epochs. Each output dataset had a duration of 100 seconds and consisted of 64 channels (standard Biosemi montage) with a sampling frequency of 256 Hz. See **Fig S1** for examples of *clean* synthetic EEG that was used.

#### EOG and EMG artifacts

In order to make the synthetic artifacts as realistic as possible, we used the real-world recordings of sensor data directly overlying the eyes (EOG, electro-oculogram) and facial muscles (EMG, electromyogram) from the EEGdenoiseNet dataset https://github.com/ncclabsustech/EEGdenoiseNet (H. Zhang et al., 2021). Then, as shown in **Figure 2**, these EOG/EMG single time-series (“waveforms”) were multiplied by dipolar source patterns (“topographies”) to generate multi-channel EEG in sensor-space. In order for these topographies to be consistent with EOG/EMG generation, we utilized the HArtMuT head model (Harmening et al., 2022), which enables simulating EEG activity originating from extra-cerebral sources located in the eyes and muscles. Here, we extracted a random sample of 16 eye and 10 scalp muscle source locations (as illustrated in **Figure S3**). Then, for each dataset, a minimum of 2 sources of EOG and 1-3 sources of EMG were randomly selected and their topographic information (i.e. source location and orientation) was retained. Finally, these EOG/EMG topographies were respectively multiplied by the EOG/EMG single time-series from EEGdenoiseNet (with 50 randomly sampled epochs of 2 seconds) to construct multi- channel datasets containing only artifacts. All EOG signals were band-pass filtered between 0.3 and 10 Hz. EMG signals were band-pass filtered between 1 to 120 Hz and notched at the powerline frequency of 50 Hz. See **Fig S2** for examples of *artifactual* semi-synthetic EEG that was used.

**Figure 2.**
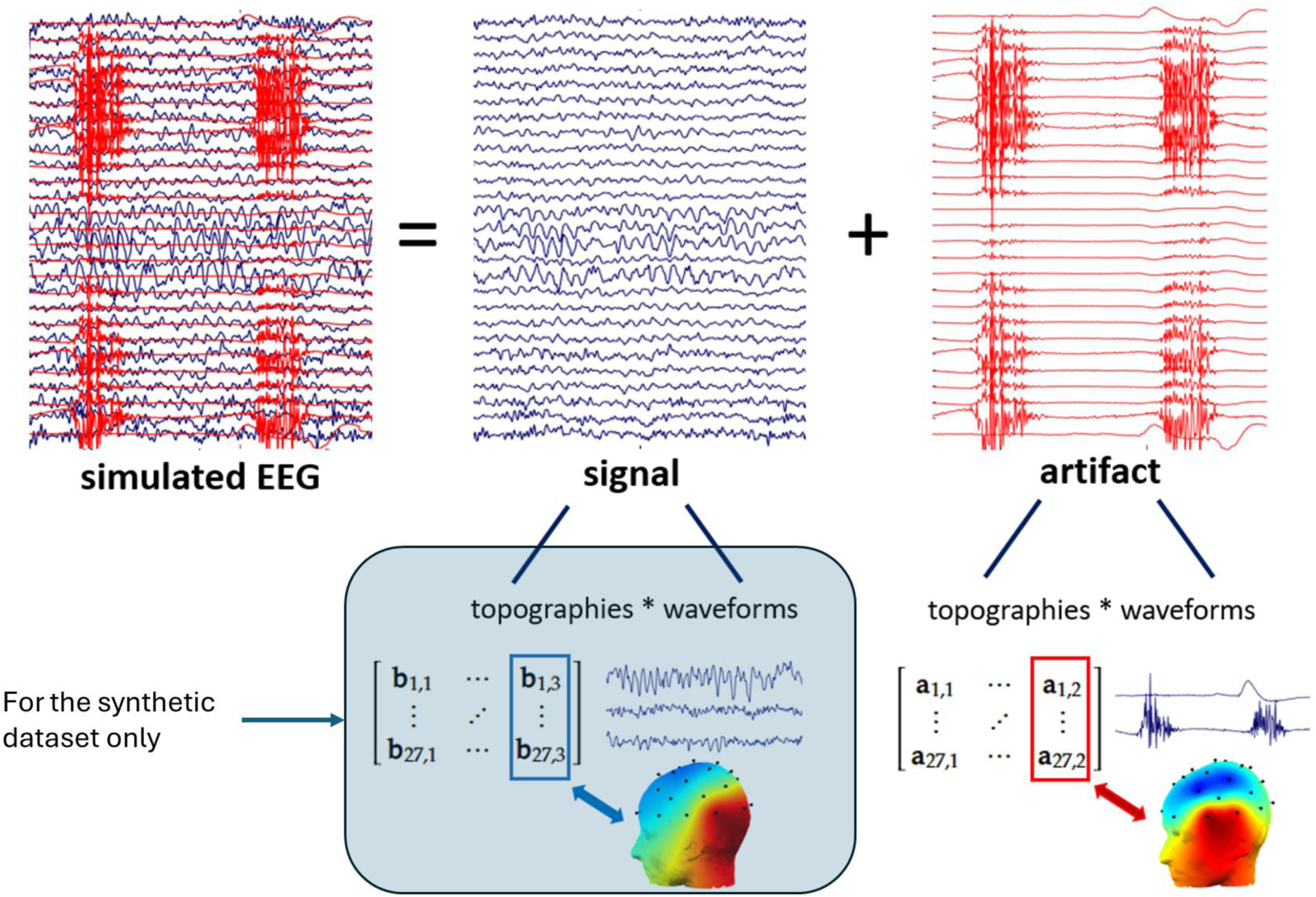
Simulated EEG data generation Background “ground-truth” EEG (signal) was obtained either from clean empirical recordings or synthetic data generated by a forward model with 100-300 sources. Contaminated (simulated EEG) data were generated by a sensor-space summation of the signal (blue trace) with different types of artifacts (red trace) (Ille et al., 2002). In this example, muscle artifact was generated by multiplying the time-series (“waveforms”) of EMG electrode activity with source locations (“topographies”) consistent with EMG artifact. Artifact waveforms were derived from the EEGdenoiseNet dataset, which consisted of direct recordings from individual EOG and EMG electrodes, respectively placed over the eyes and facial muscles (H. Zhang et al., 2021).

#### NOISE artifacts

Datasets with *noise* artifacts were identical in size to those with EOG/EMG, and were generated to contain the following types of synthetic (i.e. MATLAB simulated) “noise” across 5 datasets: with 3 *bad channels* (white noise, 1 dataset), 5 *bad channels* (sawtooth wave, 1 dataset), 6 *bad channels* (square wave, 1 dataset), *spike* artifact (synthetic, 1 dataset), impulsive noise (1 dataset).

#### Artifact mixtures

In order to challenge the denoising algorithms with more complex scenarios of artifact mixtures, the baseline EOG, EMG, and/or NOISE datasets were z-score normalised and linearly combined with each other to produce a *mixed* artifact category: “ NOISE + EMG + EOG ”. To generate a total of 5 datasets in this category, only datasets with matching numbers were added across categories (e.g. EOG dataset 1 + EMG dataset 1 + NOISE dataset 1).

In summary, 5 separate artifact datasets were generated for each of the following 4 artifact categories:

1. EMG (muscle activity)
2. EOG (eye blinks and/or movement)
3. NOISE (empirical or synthetic noise)
4. NOISE + EMG + EOG

#### Bad channels

Out of the above total of 20 artifactual datasets, there were 6 datasets containing bad channels (3 datasets in the NOISE category, and 3 datasets in the NOISE + EMG + EOG category). Thus, bad channels were present in 30% (6/20) of all synthetic simulations.

#### Mixed clean EEG + artifact signal generation

In line with the model shown in **Figure 2**, each of the 10 clean recordings was pairwise “mixed”, through linear summation, with each of 5 datasets from a specific artifact category. This resulted in 50 contaminated datasets within each one of the 4 artifact categories (EMG; EOG; NOISE; and EMG + EOG + NOISE).

Thus, the *synthetic* simulations consisted of 200 unique datasets (50 datasets x 4 artifact categories), varied across 4 temporal contamination levels x 4 SNR levels, yielding a **“synthetic” benchmark sample of 3,200 datasets**. Each dataset had 64 channels and a length of 100 seconds, sampled at 256 Hz (i.e. each data matrix: 64 channels x 25,600 samples).

### Empirical EEG simulations

#### Clean EEG signal generation

Here, we used Stefan Ehrlich’s experimentally-acquired recordings of clean EEG https://github.com/stefan-ehrlich/dataset-automaticArtifactRemoval (Ehrlich, 2019/2024), where 10 separate subjects were selected from the Ehrlich dataset in the ‘eyes-closed’ resting condition with a duration of 100 seconds. These recordings were visually inspected for the presence of any artifacts, and any samples that were suspected of contamination were wholesale rejected (i.e. removed across all channels). For every subject, clean EEG segments of at least 2 seconds were concatenated and low-pass filtered at 60 Hz. Each output dataset had a duration of 60 seconds and consisted of 27 channels with a sampling frequency of 200 Hz. See **Fig S4** for examples of *clean* empirical EEG that was used.

#### Artifact signal generation EOG and EMG artifacts

Even dedicated scalp EEG recordings of extra-cerebral artifacts inevitably contain brain signals. Consequently, ground-truth brain and artifact signals cannot be established, which is necessary for reliably evaluating denoising performance. To address this problem, we used empirical recordings of sensor data directly overlying the eyes (EOG, electro-oculogram) and facial muscles (EMG, electromyogram) from the EEGdenoiseNet dataset https://github.com/ncclabsustech/EEGdenoiseNet (H. Zhang et al., 2021). Then, as shown in **Figure 2**, the EOG/EMG single time-series (“waveforms”) can be multiplied by their relevant spatial patterns (“topographies”) to generate multi-channel EEG in sensor-space. For these topographies to be consistent with EOG/EMG generation, we separately performed (Infomax- extended) ICA on multichannel EOG/EMG artifact recordings from the Ehrlich dataset (Ehrlich, 2019/2024). Here, for each dataset, a minimum of 2 components of EOG and 4 components of EMG were manually selected and only their topographic information was retained. Finally, as described above, these EOG/EMG topographies were respectively multiplied by the EOG/EMG single time-series from EEGdenoiseNet (with 30 randomly sampled epochs of 2 seconds) to construct multi-channel datasets containing only artifacts. The use of ICA in generating the artifactual EEG data could raise concerns about circularity (i.e. "double dipping”), potentially favouring ICA-based algorithms (IClabel, MARA). Such a bias would indeed be possible if we had used the independent, non-Gaussian time courses generated by ICA itself (i.e. the time-courses matching the Ehrlich dataset topographies). However, we circumvented this issue by instead using direct EOG/EMG sensor recordings from the EEGdenoiseNet dataset as the artifact time courses. These physiological signals were obtained without imposing ICA’s assumptions of independence or non-Gaussianity, thereby ensuring a fairer evaluation of the ICA-based algorithms. All EOG signals were band-pass filtered between 0.3 and 10 Hz, and then re-sampled to 200 Hz. EMG signals were band-pass filtered between 1 to 120 Hz and notched at the powerline frequency of 50 Hz, and then resampled to 200 Hz. See **Fig S5** for examples of *artifactual* empirical EEG that was used.

#### NOISE artifacts

Datasets with *noise* artifacts were identical in size to those with EOG/EMG, and were generated to contain the following types of empirical (e.g. based on Stefan Ehrlich’s empirical recordings) and synthetic (i.e. MATLAB simulated) “noise” across 10 datasets: with 3-7 *bad channels* (synthetic, 4 datasets), *movement* artifact (empirical, 1 dataset), *electrode pop* (empirical, 1 dataset), *step* artifacts (synthetic, 1 dataset), *temporally* non-stationary noise (synthetic, 1 dataset), and *spatially* non-stationary noise (synthetic, 2 datasets). See **Fig S6** for examples of *noisy* empirical EEG that was used.

#### Artifact mixtures

In order to challenge the denoising algorithms with more complex scenarios of artifact mixtures, the baseline EOG, EMG, and/or NOISE datasets were z-score normalised and linearly combined to produce a *mixed* artifact category: “NOISE + EMG + EOG ”. To generate a total of 10 datasets in this category, only datasets with matching numbers were added across categories (e.g. EOG dataset 1 + EMG dataset 1 + NOISE dataset 1).

In summary, 10 separate artifact datasets were generated for each of the following 4 artifact categories:

1. EMG (muscle activity)
2. EOG (eye blinks and/or movement)
3. NOISE (empirical or synthetic noise)
4. NOISE + EMG + EOG

#### Bad channels

Out of the above total of 40 purely artifactual datasets, there were 8 datasets containing bad channels (4 datasets in the NOISE category, and 4 datasets in the NOISE + EMG + EOG category). Thus, bad channels were present in 20% (8/40) of all empirical simulations.

#### Mixed clean EEG + artifact signal generation

For the final step, and according to the model shown in **Figure 2** above, each of the 10 clean recordings was pairwise “mixed”, through linear summation, with each of 10 datasets from a specific artifact category. This resulted in 100 contaminated datasets within each one of the 4 artifact categories (EMG; EOG; NOISE; and EMG + EOG + NOISE).

Thus, the empirical simulations consisted of 400 unique datasets (100 datasets x 4 artifact categories), varied across 4 temporal contamination levels x 4 SNR levels, yielding an “**empirical” benchmark sample of 6,400 datasets**. Each dataset had 27 channels and a length of 60 seconds, sampled at 200 Hz (i.e. each data matrix: 27 channels x 12,000 samples).

#### Measures of denoising performance: SNR, relative root mean square error (RRMSE) and correlation coefficient

Here, clean EEG recordings served as the ground truth. Denoising algorithms were evaluated on their accuracy in recovering this signal from contaminated data using 3 standard metrics, allowing comparison with prior work (Mumtaz et al., 2021).

- ● *SNR:* higher is better

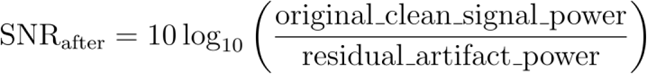

*where: original_clean_signal_power* is the variance of the original clean signal elements:

*residual_artifact_power* is the variance of the error signal (difference between denoised and original clean signal elements):

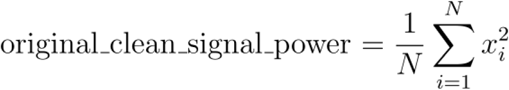

- ● *Relative Root Mean Square Error (RRMSE):* lower is better

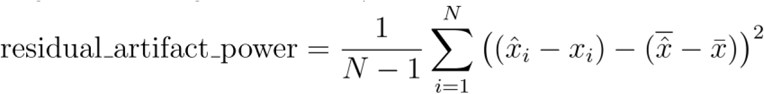

where: RMSE is the standard deviation of the error signal (difference between denoised and original clean signal elements) and RMS is the amplitude of the original clean signal elements.

- ● *Correlation Coefficient (R):* higher is bette**r**

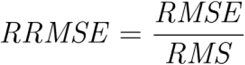

where: RMSE is the standard deviation of the error signal (difference between denoised and original clean signal elements) and RMS is the amplitude of the original clean signal elements.

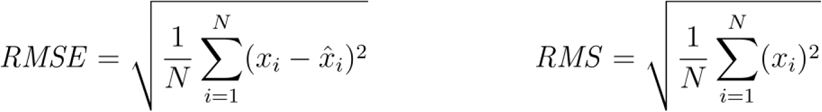

Correlation Coefficient (R): higher is better

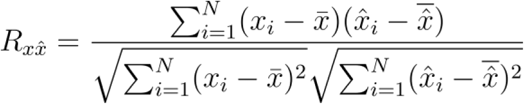

*where:*

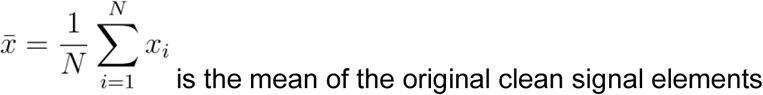

and

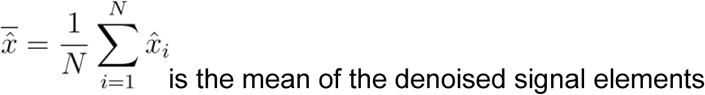

#### Statistical Analysis

To compare the benchmark performance of GEDAI against the other algorithms (ASR, ICLabel, MARA) paired, non-parametric, statistical tests were employed due to a) the same dataset(s) were denoised by all algorithms, and b) the potential non-normal distribution of performance metrics. The primary performance metrics evaluated were the Correlation Coefficient R, SNR, and RRMSE) between the denoised signal and the ground-truth clean signal. Statistical significance was set at an alpha level of 0.05. Here, we used a Friedman test (i.e. a non-parametric equivalent of a repeated-measures ANOVA) to assess if there was a statistically significant difference in the distributions of denoising performance scores between the algorithms (GEDAI, ASR, ICLabel, MARA) when applied to the *same* contaminated datasets (varied by SNRbefore, temporal contamination, and artifact type). Following a significant two-sided Friedman test result (i.e. p < 0.05) on SNRafter, post-hoc pairwise two-tailed comparisons between algorithms were conducted using MATLAB’s *multcompare* function with a Bonferroni multiple-comparison correction to control for family-wise error rate. Effect sizes for each pairwise comparison were reported using the *standardized Z-score effect size* (*r* = Z/√N), where Z is the Z-score from the Wilcoxon test and N is the number of pairs (where r ≈ 0.1 is small, r ≈ 0.3 is medium, and r ≈ 0.5 is large). All statistical analyses were performed using MATLAB (v2024b).

### Section III: Neurobehavioral Prediction

This section investigated GEDAI’s denoising performance using real-world EEG data, moving beyond simulated environments. While genuine ground-truth EEG data is unobtainable in such circumstances, EEG can still be used to predict specific objective information from an experiment. Our central hypothesis was that superior denoising would preserve the crucial neural information necessary for more accurate predictions. We tested this hypothesis across two distinct datasets and scenarios below.

#### Classification of visual stimuli based on single-trial Event Related Potentials (ERP)

Here, we used publicly available data (https://zenodo.org/records/7495536) from a visual oddball study by Omejc and colleagues (Omejc et al., 2023). A total of 70 healthy adult individuals passively viewed frequent (white square) and rare (Einstein’s face) stimuli while being recorded with a 32-channel EEG sampled at 256 Hz. Each participant was presented with 124 frequent (84%) and 23 rare (16%) stimuli. For full details of the experiment please see (Omejc et al., 2023). After 1-40 Hz band-pass filtering and average referencing the continuous EEG data, we ran each denoising algorithm with its default settings (as described in the EEG simulations) *without* any bad epoch rejection. Accordingly, ASR’s bad channel rejection step was enabled for this algorithm, as well as prior to running IClabel or MARA. No explicit bad channel rejection was utilized for GEDAI. For IClabel, only components flagged as "brain" or "other" were retained as clean EEG data. The open MATLAB analysis code (https://github.com/NinaOmejc/VEP_classification_aging) was then utilized for the extraction of single-trial ERP features (Omejc et al., 2023). Time-independent statistical ERP features were extracted based on four ERP components: P1, N170, P2, and P3. Each of these components was parameterized by four metrics: peak amplitude, mean amplitude, peak latency, and fractional 50% peak latency. After feature selection based on mutual information, eight time-independent features were used for classification, specifically the peak amplitudes and fractional 50% peak latencies at four electrode clusters (occipital, parietal, central, and frontal).

We performed the binary classification task by concatenating all subjects’ trials together, and used a linear support vector machine (Matlab’s fitcsvm function) with tenfold cross-validation, averaging the validation scores over 100 runs. In order to assess each algorithm’s full potential for signal recovery, we did not perform any “bad” ERP trial rejection before or after denoising. Hence, all trials were classified based on whether they contained a frequent or infrequent stimulus. Given the strong class imbalance, classification performance was reported as the area under the curve (AUC) of the receiver operating characteristic (ROC), averaged across all subjects. DeLong’s test (DeLong et al., 1988) was used to estimate the statistical significance of AUC between models, with Bonferroni multiple comparison correction.

#### Subject identification based on brain fingerprinting

Here, we used each person’s unique resting-state EEG activity to identify them among a group of 100 individuals, also known as “brain fingerprinting”. For this, we utilised the *Dortmund Vital Study*dataset, freely accessible at OpenNeuro (https://doi.org/10.18112/openneuro.ds005385.v1.0.3). This dataset contained EEG recordings of adult subjects (mean age: 44 years) in a test-retest design, where the retest of the same individuals was recorded at 5 years follow-up. From this dataset, we included the first n=100 subjects that contained recordings of both Session 1 (“test”) and Session 2 (“retest “). Each test and retest file consisted of 180 seconds of *eyes-open* resting state before a cognitive task, recorded with 62 channels and a sampling rate of 100 Hz. After 1-45 Hz band- pass filtering and average referencing, all EEG recordings were then denoised with ASR, IClabel, MARA, or GEDAI using default settings (identical to the EEG simulations). Hence, bad channel rejection was enabled for ASR, IClabel and MARA, but not GEDAI. For IClabel, only components flagged as "brain" or "other" were retained as clean EEG data. In order to conduct fingerprinting, we first calculated the spatial covariance matrix (i.e. channel x channel) of the denoised data using MATLAB’s *cov* function. Then, the *correlation distance* (with range -1 to +1) between the Session 1 recordings’ and Session 2 recordings’ covariance matrices was calculated using MATLAB’s *pdist2* function. The values from this square distance matrix were subtracted from 1, yielding a 100 x 100 similarity matrix between Sessions 1 (test) and Sessions 2 (retest), also known as the Identifiability matrix (Amico & Goñi, 2018). The Identifiability matrix has subjects as rows and columns, and encodes the information about the self-similarity (I*self*, main diagonal elements) of each subject with themself, across the test/retest sessions, and the similarity of each subject with the others (or I*others*, off-diagonal elements). The overall goal is to predict the identity of a single subject from Session 1 based on the similarity of that recording with multi-subject data from Session 2. As a measure of fingerprinting performance, we used the *success rate* (Sorrentino et al., 2021) from code available at https://github.com/eamico/Clinical_fingerprinting/blob/master/FC_fingerprint.m. The success rate (%) indicates how many times an I*self* value is higher than the I*others* values on the same row and column of the Identifiability matrix. In other words, the *success rate* represents the percentage of the number of individual off-diagonal comparisons when the self-correlation “wins”, which is a more granular comparison rather than a binary correct/incorrect prediction of the subject. To estimate the variability in fingerprinting performance for each algorithm, we performed 50 random resamples of the denoised EEG data. Each resample consisted of a total of 10 seconds (i.e. 1000 random data samples), from which the spatial covariance matrix was calculated and the success score estimated. Following a significant Kuskall-Wallis test result (p <0.05), post-hoc comparisons between algorithms were conducted using MATLAB’s *multcompare* function with a Bonferroni correction to control the family-wise error rate.

#### Hardware and Software Platform

All computations were carried out using MATLAB software v2024b running on a Windows desktop PC with an Intel-Core Ultra 9 285K CPU with 24 parallel cores and 64GB of RAM. All tests were performed with the CPU using parallel processing, with no GPU acceleration.

## Results

Here, we compared the GEDAI algorithm to three established denoising algorithms (ASR, MARA and ICLabel) in terms of ground-truth (i.e. clean background signal) recovery. We principally report the SNRafter and RRMSE effect sizes. Matching figures for RRMSE can be found in Fig. SX of the Supplementary Results. For the Correlation Coefficient R, we show its values in the global comparisons at the end of each section.

### Denoising Benchmark of Synthetic EEG

We compared the denoising of 3,200 contaminated datasets, each containing 100 seconds of 64-channel EEG, which varied across 4 levels of temporal contamination, baseline SNRs, and artifact type. These datasets combined synthetic background EEG (simulated *in silico* via forward modelling) with semi-synthetic EOG/EMG artifacts and synthetic noise (see **Methods** for more details).

For **example videos** of *synthetic* ground-truth, noise-contaminated and GEDAI-denoised EEG, see **Synthetic_Video_1** and **Synthetic_Video_2**.

Figure 3 above shows benchmarking across different aggregation levels (temporal contamination levels, SNRbefore, artifact type and computational time). Higher SNRafter and lower RRMSE (for RRMSE see Supplementary **Fig. S7**) values indicate better denoising, respectively. Differences in SNRafter and RRMSE are reported below (effect size *r* = Z/√N).

**Figure 3.**
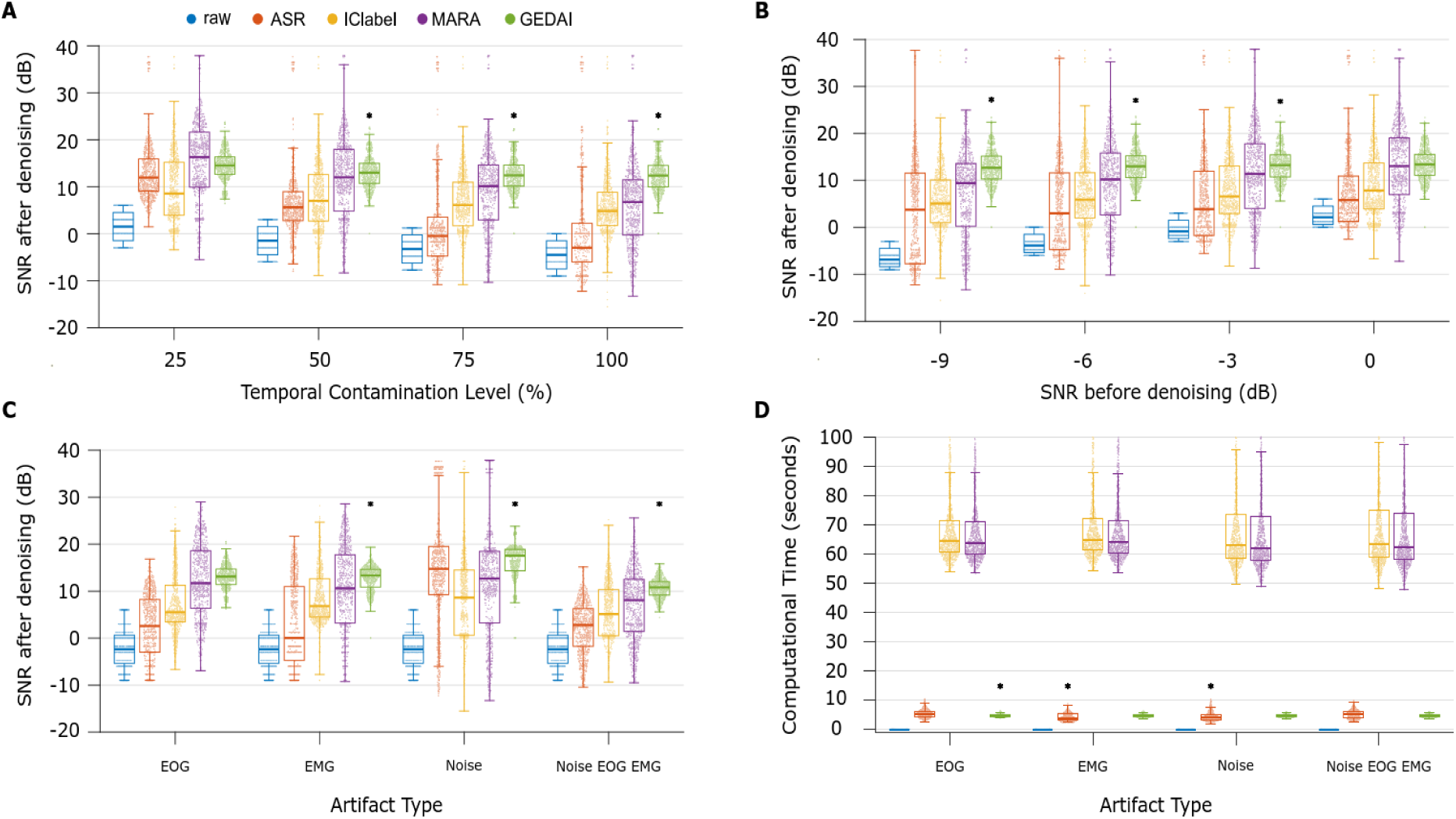
Denoising of Synthetic EEG A) By temporal contamination (%). Results are pooled across all SNRbefore and all artifact types. For every temporal contamination level, each coloured data point represents 1 of 800 datasets (50 datasets x 4 SNR levels x 4 artifact types). B) By SNRbefore (dB). Results are pooled across all temporal contamination levels and all artifact types. For every SNRbefore, each coloured data point represents 1 of 800 datasets (50 datasets x 4 artifact types x 4 temporal contamination levels). C) By artifact type. Results are pooled across all SNRbefore and temporal contamination levels. For each artifact type, each coloured data point represents 1 of 800 datasets (50 datasets x 4 SNR levels x 4 temporal contamination levels). D) Computational time (seconds). Lower values indicate faster calculations. An asterisk indicates the winning algorithm (p < 0.05 Bonferroni corrected). If no asterisk is present, it signifies a statistical tie.

#### By temporal contamination (Fig 3A)

**For 25%**, GEDAI (median = 14.54 dB, IQR = 12.67− 16.35) outperformed all algorithms *except for* MARA (median = 16.34 dB, IQR = 9.91−21.63), with non-significant differences in SNRafter (r = -0.13, p = 0.24) and RRMSE (r = 0.0082, p = 0.46); following a Friedman test: χ²(3) = 257, *p* = 2.6 x 10^-55^.

**For 50%**, GEDAI (median = 13.00 dB, IQR = 10.71−14.99) outperformed all algorithms, including the next-best MARA (median = 12.00 dB, IQR = 4.82−17.95) in SNRafter (r = 0.090, p = 0.0036) and RRMSE (r = -0.22, p = 0.0015); following a Friedman test: χ²(3) = 519, *p* = 2.9 x 10^-112^.

**For 75%**, GEDAI (median = 12.43 dB, IQR = 10.12−14.63) outperformed all algorithms, including the next-best MARA (median = 10.12 dB, IQR = 2.93−14.65) in SNRafter (r = 0.34, p = 6.4e-15) and RRMSE (r = -0.46, p = 1.9 x 10^-18^); following a Friedman test: χ²(3) = 899, *p* = 1.50 x 10^-194^.

**For 100%**, GEDAI (median = 12.38 dB, IQR = 9.96−14.55) outperformed all algorithms, including the next-best MARA (median = 6.75 dB, IQR = -0.28−11.47), the closest competitor, in SNRafter (r = 0.59, p = 4.7 x 10^-51^) and RRMSE (r = -0.70, p = 1.8 x 10^-60^); following a Friedman test: χ²(3) = 1028, *p* = 1.6 x 10^-222^.

#### By SNRbefore (**Fig 3B**)

**For -9 dB**, GEDAI (median = 12.68 dB, IQR = 10.28−15.16) outperformed all algorithms, including the next-best MARA (median = 9.39 dB, IQR = 0.23−13.59), the closest competitor, in SNRafter (r = 0.46, p = 1.40 x 10^-30^) and RRMSE (r = -0.56, p = 3.7 x 10^-33^); following a Friedman test: χ²(3) = 657, *p* = 4.3 x 10^-142^.

**For -6 dB**, GEDAI (median = 13.00 dB, IQR = 10.59−15.24) outperformed all algorithms, including the next-best MARA (median = 10.19 dB, IQR = 2.63− 15.81), the closest competitor, in SNRafter (r = 0.33, p = 2.0 x 10^-16^) and RRMSE (r = -0.46, p = 1.3 x 10^-19^); following a Friedman test: χ²(3) = 607, *p* = 2.4 x 10^-131^.

**For -3 dB**, GEDAI (median = 13.23 dB, IQR = 10.72−15.43) outperformed all algorithms, including the next-best MARA (median = 11.36 dB, IQR = 4.01−17.78), the closest competitor, in SNRafter (r = 0.14, p = 0.0025) and RRMSE (r = -0.30, p = 3.7 x 10^-05^); following a Friedman test: χ²(3) = 565, *p* = 3.2 x 10^-122^.

**For 0 dB**, GEDAI (median = 13.37 dB, IQR = 11.06−15.52) outperformed all algorithms *except for* MARA (median = 13.04 dB, IQR = 6.97−19.05), with non-significant differences in SNRafter (r = -0.03, p = 1) and RRMSE (r = -0.14, p = 0.83); following a Friedman test: χ²(3) = 493, *p* = 1.8 x 10^-106^.

#### By artifact type (Fig 3C)

**For EOG**, GEDAI (median = 13.12 dB, IQR = 11.41−14.71) outperformed all algorithms *except for* MARA (median = 11.70 dB, IQR = 6.35−18.62), with non-significant differences in SNRafter (r = 0.049, p = 0.79) and RRMSE (r = -0.16, p = 0.79); following a Friedman test: χ²(3) = 1127, *p* = 4.1 x 10^-244^.

**For EMG**, GEDAI (median = 13.33 dB, IQR = 10.87−14.65) outperformed all algorithms, including the next-best MARA (median = 10.62 dB, IQR = 3.22−17.75) in SNRafter (r = 0.25, p = 1.9 x 10^-10^) and RRMSE (r = -0.36, p = 1.6 x 10^-10^); following a Friedman test: χ²(3) = 614, *p* = 1.0 x 10^-132^.

**For NOISE**, GEDAI (median = 17.54 dB, IQR = 14.35 - 18.88) outperformed all algorithms, including the next-best ASR (median = 14.75 dB, IQR = 9.28−19.51) in SNRafter (r = 0.073, p = 3.8 x 10^-06^) and RRMSE (r = -0.33, p = 5.2 x 10^-11^); following a Friedman test: χ²(3) = 315, *p* = 5.9 x 10^-68^.

**For NOISE + EOG + EMG**, GEDAI (median = 10.82 dB, IQR = 9.16−12.02) outperformed all algorithms, including the next-best MARA (median = 8.10 dB, IQR = 1.43−12.50) in SNRafter (r = 0.44, p = 3.4 x 10^-21^) and RRMSE (r = -0.52, p = 4.7 x 10^-24^); following a Friedman test: χ²(3) = 942, *p* = 7.7 x 10^-204^.

#### Computational time (Fig 3D)

**For EOG**, GEDAI (median = 4.62 s, IQR = 4.38−4.97) outperformed all algorithms, including the next-best ASR (median = 5.30 seconds, IQR = 4.33 - 6.16) in Time (*r* = -0.44, p = 5.1 x 10^-05^); following a Friedman test: χ²(3) = 2180, *p* < m.p.

**For EMG**, GEDAI (median = 4.61 s, IQR = 4.34−4.96) outperformed all algorithms *except for* ASR (median = 3.77 seconds, IQR = 3.10−5.18), with a significant difference in Time (r = 0.40, p = 2.1 x 10^-9^); following a Friedman test: χ²(3) = 2199, *p* < m.p.

**For NOISE**, GEDAI (median = 4.48 s, IQR = 4.26−4.82) outperformed all algorithms *except for* ASR (median = 4.00 s, IQR = 3.36−4.97), with a significant difference in Time (r = 0.24, p = 1.6 x 10^-9^); following a Friedman test: χ²(3) =2200, *p* < m.p.

**For NOISE + EOG + EMG**, GEDAI (median = 4.52 s, IQR = 4.27−4.89) outperformed all algorithms *except for* ASR (median = 5.04, IQR = 3.83−5.99), with a non-significant difference in Time (r = -0.29, p = 0.13); following a Friedman test: χ²(3) = 2165, *p* < m.p.

**Globally:** following a Friedman test χ²(3) = 2252, *p* < m.p. across all datasets and conditions, GEDAI outperformed all algorithms in post-hoc pairwise comparisons, with absolute means and 95% confidence intervals illustrated in Fig. 4. GEDAI significantly differed from MARA, the closest competitor in SNRafter (r = 0.23 ; GEDAI mean = 13.1 dB ; MARA mean = 10.8 dB), RRMSE (r = -0.37 ; GEDAI mean = 0.24 ; MARA mean = 0.51) and Correlation Coefficient (r

**Figure 4.**
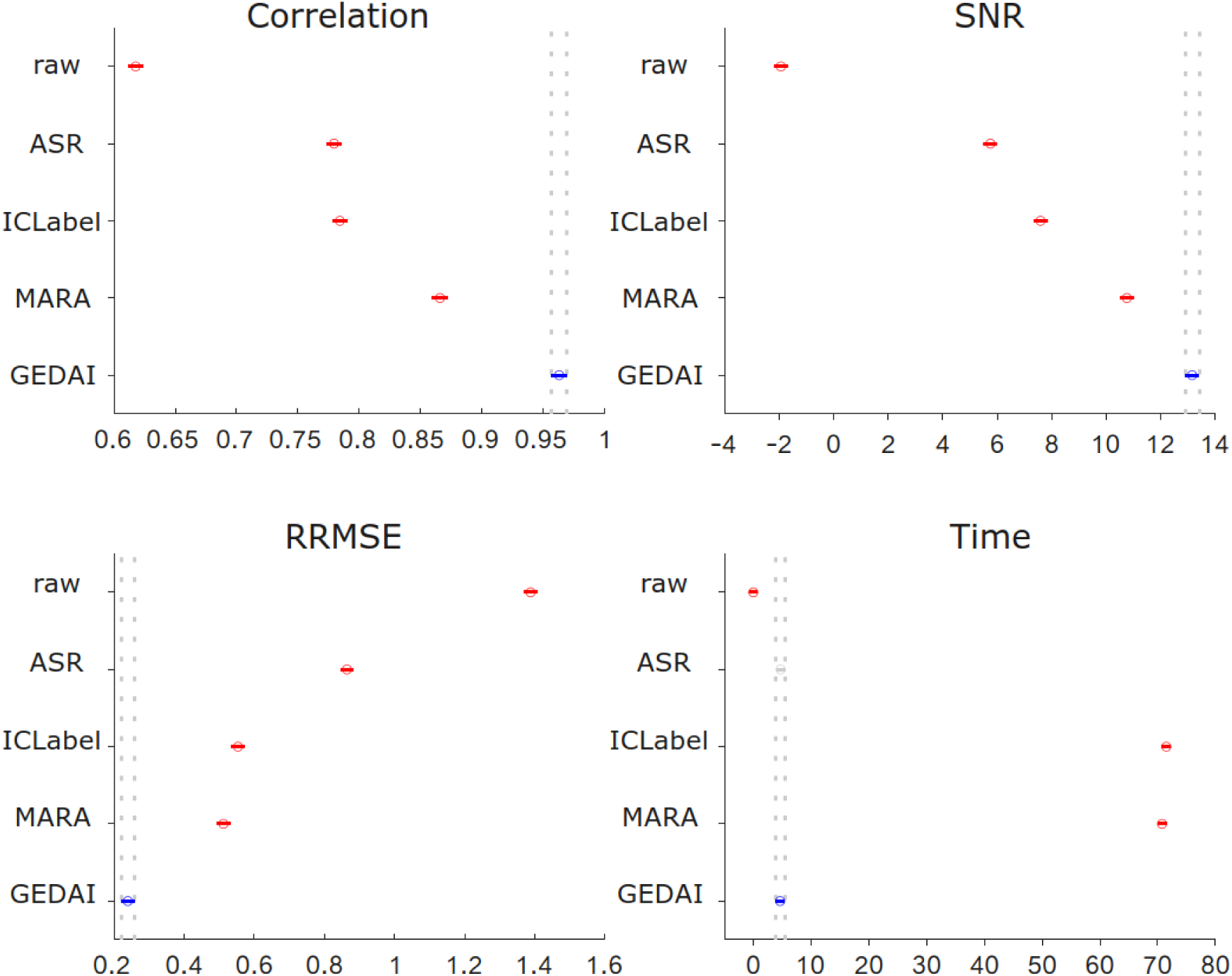
Global statistics for synthetic EEG benchmark Global means and 95% confidence intervals for synthetic EEG denoising across all temporal contamination levels, baseline SNRs and artifact types for: Correlation (top left panel, higher is better), SNRafter in dB (top right panel, higher is better), RRMSE (bottom left panel, lower is better), Computational time in seconds (bottom right panel, lower is better). Confidence intervals are plotted but are very narrow due to the large sample size.

= 0.37 ; GEDAI mean = 0.96 ; MARA mean = 0.87). GEDAI did not significantly differ from ASR, the closest competitor in Time (r = -0.03 ; GEDAI mean = 4.6 s; ASR mean = 4.7 s).

### Denoising Benchmark of Empirical EEG

We contrasted the denoising of 6,400 contaminated datasets, each containing 60 seconds of 27-channel EEG, which varied across 4 levels of temporal contamination, baseline SNR (SNRbefore), and artifact type. These datasets combined genuine background EEG (serving as the ground truth) with actual EOG/EMG/NOISE artifacts (see **Methods** for more details).

For **example videos** of *empirical* ground-truth, noise-contaminated and GEDAI-denoised EEG, see **Empirical_Video_1** and **Empirical_Video_2**.

Figure 5 above shows benchmarking across different aggregation levels (temporal contamination levels, SNRbefore, artifact type and computational time). Higher SNRafter and lower RRMSE (for RRMSE see Supplementary **Fig. S7**) values indicate better denoising, respectively. Differences in SNRafter and RRMSE are reported below (effect size *r* = Z/√N).

**Figure 5.**
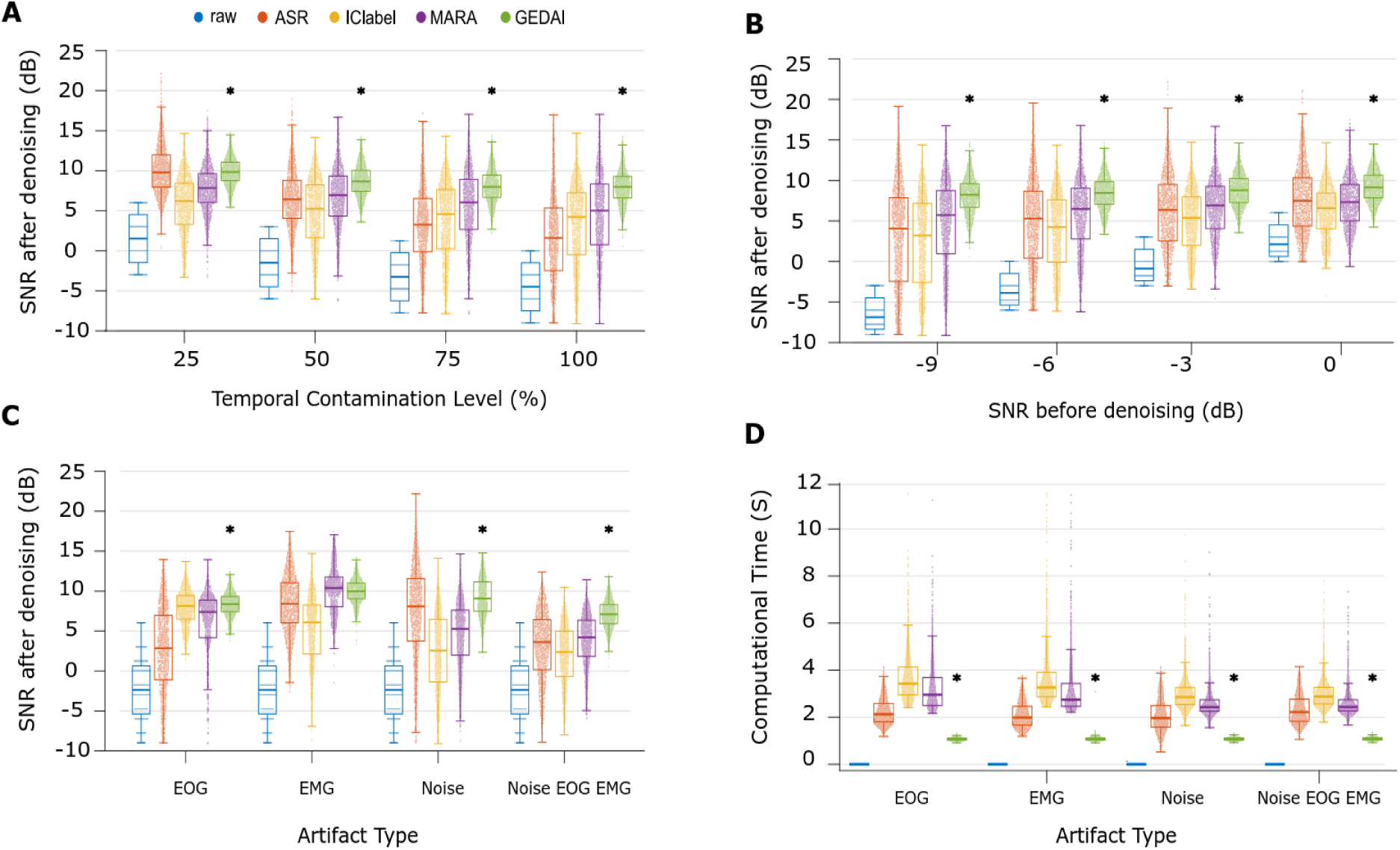
Denoising of Empirical EEG A) By temporal contamination (%). Results are pooled across all SNRbefore and all artifact types. For every temporal contamination level, each coloured data point represents 1 of 1,600 datasets (100 datasets x 4 SNR levels x 4 artifact types). B) By SNRbefore (dB). Results are pooled across all temporal contamination levels and all artifact types. For every SNRbefore, each coloured data point represents 1 of 1,600 datasets (100 datasets x 4 artifact types x 4 temporal contamination levels). C) By artifact type. Results are pooled across all SNRbefore and temporal contamination levels. For each artifact type, each coloured data point represents 1 of 1,600 datasets (100 datasets x 4 SNR levels x 4 temporal contamination levels). D) Computational time (seconds). Lower values indicate faster calculations. An asterisk indicates the winning algorithm (p < 0.05 Bonferroni corrected). If no asterisk is present, it signifies a statistical tie .

#### By temporal contamination (Fig 5A)

**For 25%**, GEDAI (median = 9.84 dB, IQR = 8.77-11.06) outperformed all algorithms, including the next-best ASR (median = 9.79 dB, IQR = 7.97-11.97) in SNRafter (r = -0.07, p = 0.032) and RRMSE (r = 0.0046, p = 0.02); following a Friedman test: χ²(3) = 1855, *p* < m.p.

**For 50%**, GEDAI (median = 8.66 dB, IQR = 7.43-10.03) outperformed all algorithms, including the next-best MARA (median = 6.95 dB, IQR = 4.34-9.33) in SNRafter (r = 0.55, p = 4.8 x 10^-90^) and RRMSE (r = -0.60, p = 2.2 x 10^-91^); following a Friedman test: χ²(3) = 1134, *p* = 1.6 x 10^-^245.

**For 75%**, GEDAI (median = 7.98 dB, IQR = 6.67-9.45) outperformed all algorithms, including the next-best MARA (median = 6.06 dB, IQR = 2.67-8.92) in SNRafter (*r* = 0.53, p = 1.0 x 10^-65^) and RRMSE (*r =* -0.60, p = 1.1 x 10^-73^); following a Friedman test: χ²(3) = 1356, *p* = 1.3 x 10293.

**For 100%**, GEDAI (median = 7.98 dB, IQR = 6.60-9.27) outperformed all algorithms, including the next-best MARA (median = 5.01 dB, IQR = 0.76-8.34) in SNRafter (*r* = 0.62, p = 6.2 x 10^-105^) and RRMSE (*r =* -0.67, p = 2.3 x 10^-112^); following a Friedman test: χ²(3) = 1615, *p* < m.p. **by SNRbefore (Fig 5B)**

**For -9 dB**, GEDAI (median = 8.24 dB, IQR = 6.69−9.60) outperformed all algorithms, including the next-best MARA (median = 5.73 dB, IQR = 0.97−8.74) in SNRafter (r = 0.61, p = 1.92 x 10^-101^) and RRMSE (r = -0.66, p = 2.1 x 10^-104^); following a Friedman test: χ²(3) =1334, *p* = 8.3 x _10_-289.

**For -6 dB**, GEDAI (median = 8.46 dB, IQR = 7.07−9.83) outperformed all algorithms, including the next-best MARA (median = 6.49 dB, IQR = 2.77−9.03) in SNRafter (r = 0.58, p = 3.0 x 10^-91^) and RRMSE (r = -0.64, p = 1.1 x 10^-97^) ; following a Friedman test: χ²(3) =1194, *p* = 1.5 x _10_-258.

**For -3 dB**, GEDAI (median = 8.79 dB, IQR = 7.28−10.24) outperformed all algorithms, including the next-best MARA (median = 6.92 dB, IQR = 4.06−9.27) in SNRafter (r = 0.58, p = 9.8 x 10^-92^) and RRMSE (r = -0.63, p = 1.5 x 10^-98^); following a Friedman test: χ²(3) = 1078, *p* = 2.7 x 10^-233^.

**For 0 dB**, GEDAI (median = 9.15 dB, IQR = 7.85−10.62) outperformed all algorithms, including the next-best ASR (median = 7.49 dB, IQR = 4.35−10.34) in SNRafter (r = 0.47, p = 9.2 x 10^-85^) and RRMSE (r = -0.55, p = 5.9e-88); following a Friedman test: χ²(3) =1056, *p* = 1.6 x 10^-228^.

### By artifact type (Fig 5C)

**For EOG**, GEDAI (median = 8.37 dB, IQR = 7.44−9.32) outperformed all algorithms, including the next-best IClabel (median = 8.15 dB, IQR = 6.50−9.45) in SNRafter (*r* = 0.26, p = 0.003) and RRMSE (*r =* -0.30, p = 0.003); following a Friedman test: χ²(3) = 1918, *p* < m.p.

**For EMG**, GEDAI (median = 9.97 dB, IQR = 9.04−10.99) outperformed all algorithms, *except for* MARA (median = 10.39 dB, IQR = 8.04−11.76), with non-significant differences in SNRafter (r = -0.06, p = 0.70) and RRMSE (r = -0.007, p = 0.70); following a Friedman test: χ²(3) = 1574, *p* < m.p.

**For NOISE**, GEDAI (median = 9.08 dB, IQR = 7.47−11.16) outperformed all algorithms, including the next-best ASR (median = 8.08 dB, IQR = 3.76−11.56) in SNRafter(r = 0.24, p = 6.5e-17) and RRMSE (r = -0.40, p = 7.9e-26); following a Friedman test: χ²(3) =2301, *p* < m.p. **For NOISE + EOG + EMG**, GEDAI (median = 7.1 dB, IQR = 5.93−8.29) outperformed all algorithms, including the next-best MARA (median = 4.21 dB, IQR = 1.83−6.36) in SNRafter (r = 0.82, p = 8.4 x 10^-186^) and RRMSE (r = -0.83, p = 1.9 x 10^-192^); following a Friedman test: χ²(3) = 2478, *p* < m.p.

#### Computational time (Fig 5D)

**For EOG**, GEDAI (median = 1.06 s, IQR = 1.03−1.11) outperformed all algorithms, including the next-best ASR (median = 2.13 s, IQR = 1.80−2.58) in Time (*r* = -0.87, p = 6.0 x 10^-141^); following a Friedman test: χ²(3) = 4493, *p* < m.p.

**For EMG**, GEDAI (median = 1.07 s, IQR = 1.03−1.11) outperformed all algorithms, including the next-best ASR (median = 1.99 s, IQR = 1.67−2.47), the closest competitor, in Time (*r* = - 0.87, p = 4.3 x 10^-134^); following a Friedman test: χ²(3) = 4562, *p* < m.p.

**For NOISE**, GEDAI (median = 1.07 s, IQR = 1.03−1.12) outperformed all algorithms, including the next-best ASR (median = 1.96 s, IQR = 1.58−2.50) in Time (r = -0.85, p = 6.1 x 10^-161^); following a Friedman test: χ²(3) = 4098, *p* < m.p.

**For NOISE + EOG + EMG**, GEDAI (median = 1.08 s, IQR = 1.04 - 1.13) outperformed all algorithms, including the next-best ASR (median = 2.22 s, IQR = 1.83−2.77) in Time (r = - 0.87, p = 1.53 x 10^-223^); following a Friedman test: χ²(3) = 4018, *p* < m.p.

**Globally:** following a Friedman test χ²(3) = 4704, *p* < m.p. across all datasets and conditions, GEDAI outperformed all algorithms in post-hoc pairwise comparisons (*p* < m.p.), with absolute means and 95% confidence intervals illustrated in Fig. 6. GEDAI significantly differed from MARA, the closest competitor in SNRafter (r = 0.59 ; GEDAI mean= 8.67 dB ; MARA mean = 6.14 dB) and RRMSE (r = -0.64 ; GEDAI mean = 0.38 ; MARA mean = 0.57). GEDAI significantly differed from ASR, the closest competitor, in Correlation Coefficient (r = 0.63 ; GEDAI mean= 0.92 ; ASR mean= 0.83) and Time (r = -0.86 ; GEDAI mean = 1.1 s; ASR mean = 2.2 s).

**Figure 6.**
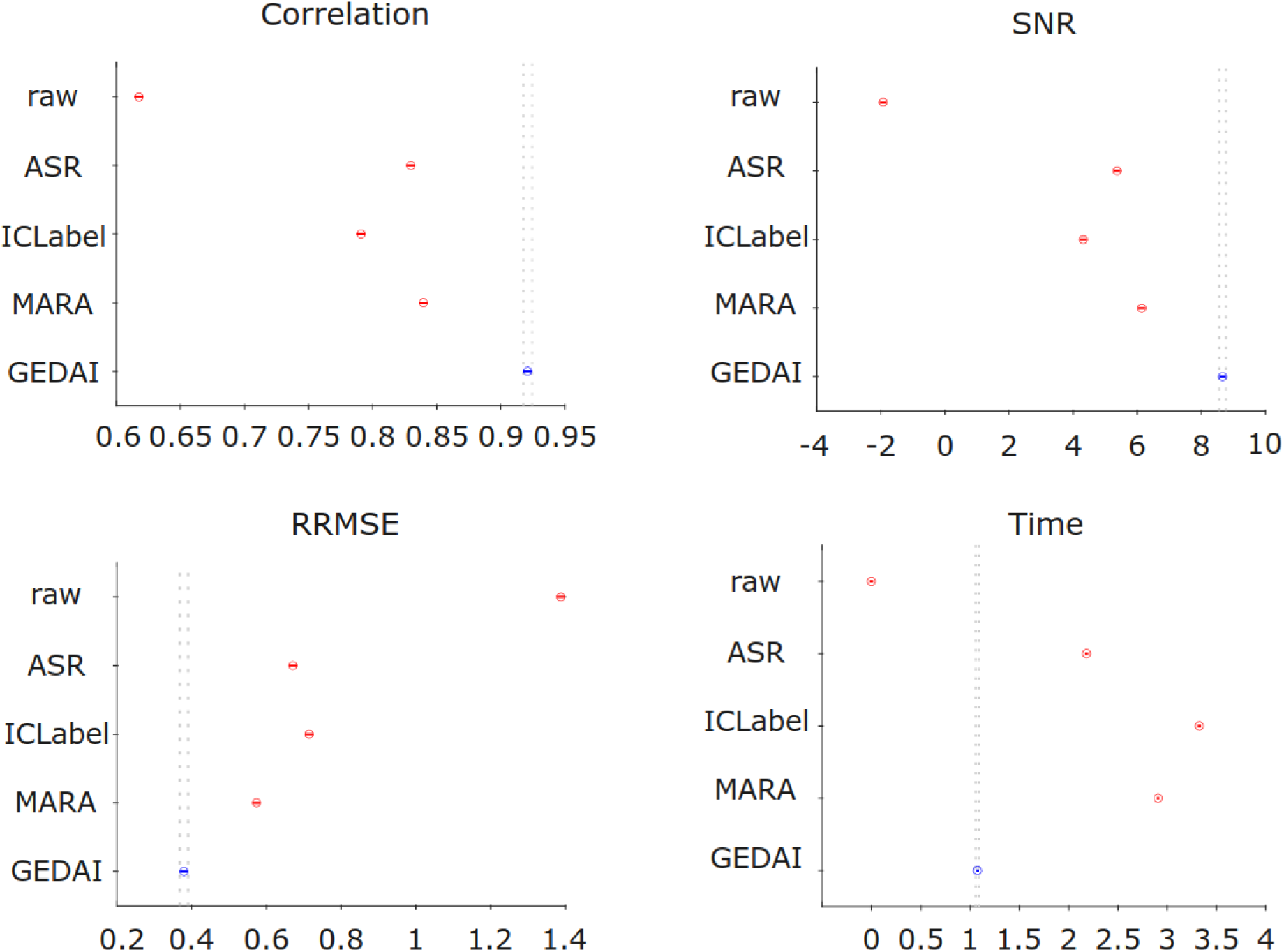
Global statistics for empirical EEG benchmark Global means and 95% confidence intervals for empirical EEG denoising across all temporal contamination levels, baseline SNRs and artifact types for: Correlation (top left panel, higher is better), SNRafter in dB (top right panel, higher is better), RRMSE (bottom left panel, lower is better), Computational time in seconds (bottom right panel, lower is better). Confidence intervals are plotted but are very narrow due to the large sample size.

### Neurobehavioral Prediction

#### Single-trial ERP Classification

The goal here was to predict whether a subject observed either a frequent (‘white square’) or infrequent (‘Einstein’s face’) visual stimulus, based on a single-trial ERP. Figure 7A shows the receiver-operating characteristics (ROCs) indicating binary classification performance following denoising by each algorithm without any bad trial rejection. DeLong tests on the area-under-the-curve (AUC) confirmed that GEDAI (mean AUC = 0.80) statistically outperformed (p = 4.2 x 10^-36^) the next-best algorithm ASR (mean AUC = 0.72), followed by MARA (mean AUC = 0.58) and IClabel (mean AUC = 0.58). Basic average referencing plus 1-40 Hz band-pass filtering, i.e RAW (mean AUC = 0.52) performed close to chance level (AUC of 0.50, dotted-line).

**Figure 7.**
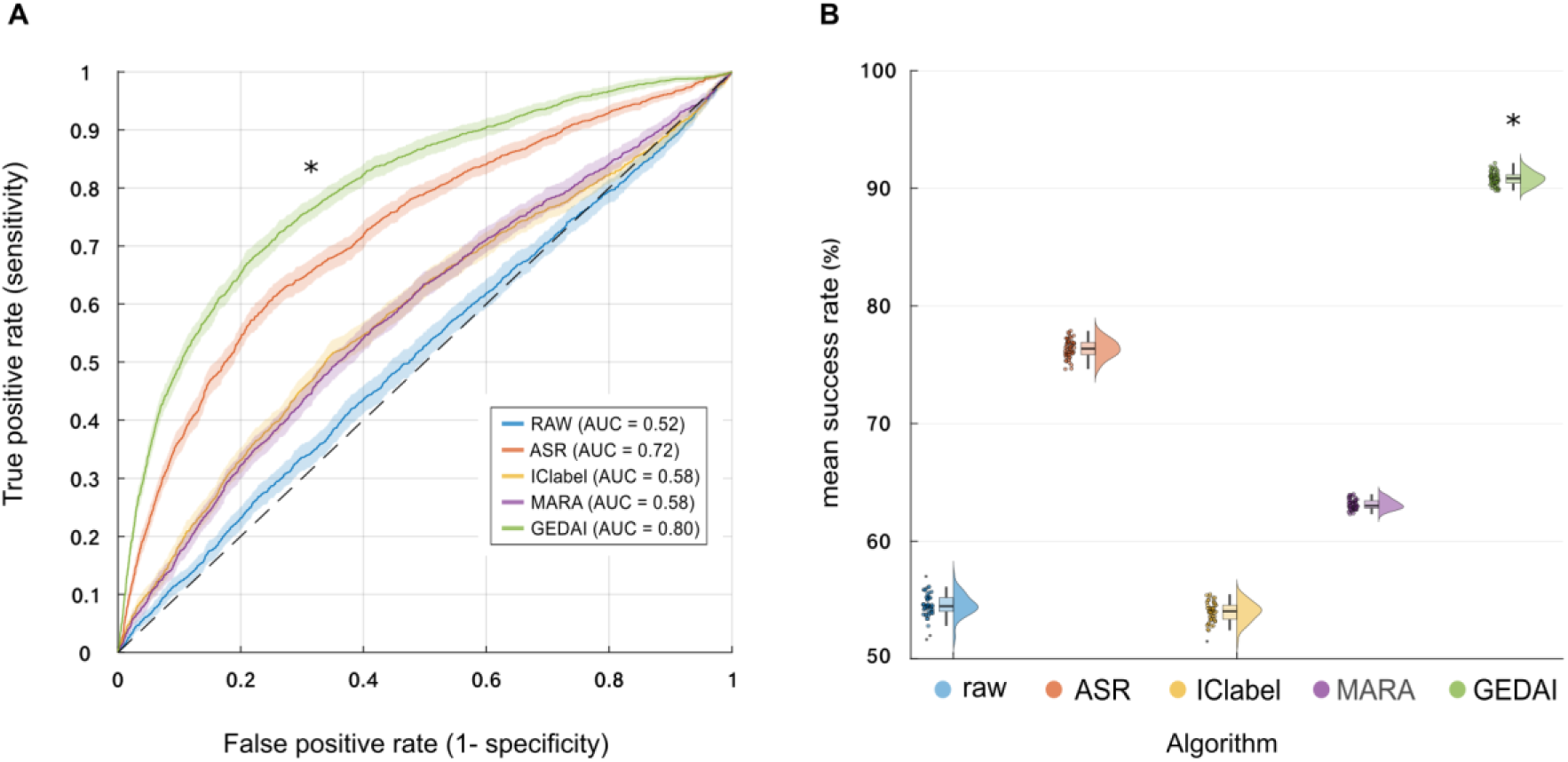
Single-trial ERP Classification and Brain Fingerprinting A) ROC curves for binary classification of ERPs from a visual oddball task, after denoising with each algorithm. Each curve is based on >10,000 trials (no bad trials were rejected). B) Mean success rate (%) of individual identification from 100 subjects, after denoising with each algorithm. Each point represents 1 of 50 resamples from each subject’s resting-state EEG data. An asterisk indicates the winning algorithm (p < 0.05 Bonferroni corrected).

#### Brain Fingerprinting

In the context of brain fingerprinting, and as shown in Figure 7B, following a significant Kuskall-Wallis test χ²(3) = 187, p = 3.4 x 10^-40^, post-hoc comparisons indicated that GEDAI (mean success rate = 91%) demonstrated higher accuracy (r = 0.86, p = 9.4 x 10^-5^) than the next-best algorithm ASR (mean success rate = 76%), followed by MARA (mean success rate = 63%), RAW (mean success rate = 55%) and IClabel (mean success rate = 54%).

Feature Comparison between Denoising Algorithms

## Discussion

This paper introduces GEDAI, a novel algorithm designed to advance EEG artifact correction, addressing a key challenge in advancing EEG technology, as highlighted in a survey by Mushtaq and colleagues (Mushtaq et al., 2024). Their study, involving 500 experts across more than 50 countries, identified improved artifact correction as the top priority in the field. Due to its theoretical underpinnings, GEDAI is fully automated, requiring no user expertise or input, and may be used to recover brain signals from low signal-to-noise recordings (e.g. -9 dB) with up to 100% temporal contamination. Validated through rigorous simulations on thousands of synthetic and real-world datasets, GEDAI marks a significant advance in EEG artifact removal, delivering a unique blend of processing speed and denoising precision. As elaborated below, GEDAI consistently performed better than, or on par with, competing algorithms in simulations containing a single type of artifact (EMG or EOG only). Interestingly, its most significant performance improvements were evident when handling multiple spatially non-stationary and temporally overlapping artifacts, such as NOISE + EMG + EOG. This makes it especially promising for use in real-world, noise-agnostic environments.

### Benchmark Simulations: synthetic and empirical EEG

Across both synthetic and empirical EEG benchmarks, GEDAI proved to be a highly effective and robust method for artifact removal, demonstrating superior performance under diverse and challenging conditions.

GEDAI’s key strength was its superior performance on heavily contaminated data. While comparable to other algorithms like ASR and MARA in datasets with sparse artifacts (25% contamination), its superiority grew significantly as contamination levels increased to 50%, 75%, and 100%, making it optimal for processing pervasively corrupted recordings.

Furthermore, GEDAI excelled at handling complex, mixed artifacts. While its performance on isolated ocular or muscular artifacts was similar to that of ICA-based methods, it showed a large effect size of |r| > 0.5 when disentangling mixtures of EOG, EMG, and noise—a crucial capability for real-world applications.

Finally, the algorithm consistently outperformed competitors in low signal-to-noise ratio (SNR) environments (from -9 dB to -3 dB). This ability to reliably recover weak neural signals from substantial background noise confirms GEDAI’s suitability for applications where data quality is a significant challenge.

### Neurobehavioral Prediction: ERP classification and brain fingerprinting

The neurobehavioral results demonstrate that the GEDAI algorithm significantly enhances the quality of EEG data for machine learning applications, outperforming all other tested methods in two distinct tasks. For single-trial ERP classification, GEDAI achieved a high mean AUC of 0.80, substantially better than the next-best algorithm, ASR (0.72), and far exceeding the near-chance performance of raw data. This performance advantage was even more pronounced in the brain fingerprinting paradigm, where GEDAI reached 91% accuracy in subject identification, surpassing ASR (76%) and other methods that were only marginally better than baseline.

### GEDAI Compared to Existing Denoising Methods

The GEDAI algorithm offers a combination of advantages that, to the best of our knowledge, is not currently exhibited by competing EEG denoising methods within a single package. As shown in Table 1, GEDAI appears to globally encompass the major strengths of ASR, ICLabel, and MARA, while their capabilities are limited to specific subsets of features. GEDAI’s favorable outcomes may be firstly attributed to the adaptability of GEVD decomposition, consistent with previous reports (Gouy-Pailler et al., 2009; Somers et al., 2018). Unlike PCA, GEVD is not limited by source orthogonality, nor is it restricted by Gaussianity, as is the case with ICA (Cohen, 2022b).

**Table 1.** Algorithmic feature comparison of the tested denoising algorithms Note: ✓ means that the performance of the function should not be affected in the listed condition. However, edge cases in which a technique could perform to a satisfying level in a condition for which it is not marked with an ✓ are possible.

| Key features of state-of-the-art denoising algorithms |  |  |  |  |
| --- | --- | --- | --- | --- |
|  | ASR | ICLabel | MARA | GEDAI |
| Rank preservation | ✓ | - | - | ✓ |
| Short data length (<1 second) | ✓ | - | - | ✓ |
| Automatic thresholding / feature selection | - | ✓ | ✓ | ✓ |
| Non-orthogonal artifacts | - | ✓ | ✓ | ✓ |
| Gaussian artifacts | ✓ | - | - | ✓ |
| Spatially non-stationary artifacts | ✓ | - | - | ✓ |
| 100% temporally contaminated data | - | ✓ | ✓ | ✓ |
| Real-time capable | ✓ | - | - | ✓ |

With parallel processing, GEDAI was between 3 to 15 times faster than ICA-based denoising, and achieved speeds comparable to PCA-based ASR, a real-time capable method (Kothe & Jung, 2015). Hence, GEDAI is computationally light enough for BCI applications.

Perhaps GEDAI’s biggest advantage is that it is fully automated and requires no operator expertise or input for component selection and/or hyperparameter tuning. While these capacities are partially shared with the other algorithms, GEDAI can achieve automation in the absence of any calibration or training data. Harnessing a "noise-free" EEG leadfield as a theoretical reference model allows GEDAI to avoid shortcomings related to the identification of clean data, as is the case for PCA, or specific artifact types, as is the case for ICA. This overall combination of features makes GEDAI a great candidate for noise-agnostic settings, such as dry-electrode or mobile EEG recordings, where clean samples might not be available or the noise could be out-of-distribution.

### Leadfield Filtering: Theoretically Informed M/EEG Artifact Removal

GEDAI’s key strength is the use of a theoretical M/EEG leadfield matrix for guiding the artifact removal, a process we refer to as ‘leadfield filtering’. Existing denoising methods that use theoretical reference models include Signal Space Projection (SSP) (Uusitalo & Ilmoniemi, 1997), Signal Space Separation (SSS) (Taulu et al., 2004), PureEEG (Hartmann et al., 2014), SPHARA (Graichen et al., 2015) and the SOUND algorithm (Mutanen et al., 2018). Of these, PureEEG and SOUND explicitly leverage leadfield modeling based on head anatomy. However, unlike SOUND, GEDAI’s algorithm does not require ill-posed inverse modeling used for source activity estimation (Mutanen et al., 2018). The PureEEG algorithm, in contrast to GEDAI’s, separately computes the artifact covariance in the frequency domain using a Bayesian estimator, with the assumption that brain and artifact components are statistically uncorrelated (Hartmann et al., 2014).

### Cutting through the Noise with SENSAI

In GEDAI, a reference covariance matrix (refCOV) models the expected spatial correlations of brain activity based on a physical head model. Using joint diagonalization (de Cheveigné & Parra, 2014), GEDAI decomposes recorded data into components. Those components with a spatial covariance inconsistent with the brain model are identified as artifacts (large eigenvalues), while consistent components are treated as neural signals (small eigenvalues). Determining the optimal cutoff between the artifact and brain components is neither obvious nor trivial. GEDAI solves this with its second key innovation: the Signal & Noise Subspace Alignment Index (SENSAI). SENSAI automatically finds the best separation threshold by calculating an alignment score for various cutoffs and selecting the one that maximizes the similarity between the denoised data and the theoretical brain subspace defined by refCOV. The resulting SENSAI score is a relative, not absolute, measure of denoised data quality (0-100%), dependent on the specific head model used. Although beyond the scope of this paper, it also has potential future use as a quantitative index for comparing EEG recording quality or the performance of different denoising methods.

### Theoretical vs. Empirical Denoising Frameworks

This subject concerns the apparent dichotomy between the *theoretical* (leadfield-driven) reference matrix used by GEDAI, and the *empirical* (data-driven) reference matrix used by existing GEVD implementations for M/EEG denoising (Haslacher et al., 2021; F. Wang et al., 2025). The core trade-off between using a leadfield-based theoretical versus an empirical *refCOV* for M/EEG denoising centers on model accuracy versus data dependency. A data-driven, empirical *refCOV* captures the actual spatial structure of noise or signal as it manifests in that specific recording and subject, but still requires that suitable segments of EEG can be reliably identified and/or denoised *a priori*. In contrast, using a theoretical *refCOV* bypasses the often difficult challenge of identifying pure noise or clean signal EEG segments, but relies on a good fit between the theoretical model and the actual data. In this case, deviations from the forward model that are specific to the subject (e.g. anatomical variations) or the recording (e.g. electrode misplacement) will likely lead to greater inaccuracies in the denoising process.

### Bad channels and how to deal with them

Traditional methods handle bad channels by rejecting them entirely (Bigdely-Shamlo et al., 2015; Kumaravel et al., 2022), which removes both artifactual and potentially useful neural signals. GEDAI offers a novel alternative by avoiding this binary rejection. It instead treats activity from compromised channels as artifactual components, which are identified via GEVD and removed if their spatial characteristics deviate from the theoretical brain signal model. Although our benchmark datasets included bad channels, this study did not explicitly evaluate GEDAI’s effectiveness in correcting them at individual-channel level. Crucially, the other algorithms benefited from a bad-channel rejection pre-processing step only when bad channels were present. This potential advantage was not extended to the fully noise-agnostic GEDAI pipeline. Future work should assess whether a hybrid approach, combining dedicated bad channel rejection with GEDAI, outperforms either method alone in specific noise scenarios.

### Potential Future Extensions of GEDAI

The simultaneous acquisition of EEG-fMRI or non-invasive brain stimulation (NIBS) techniques like transcranial direct current stimulation (tDCS), transcranial alternating current stimulation (tACS), and transcranial magnetic stimulation (TMS) present significant denoising challenges due to large, complex artifacts often contaminating the entire recording. Given that GEDAI uses leadfield filtering to separate brain activity from extra-cerebral sources of noise, it may also prove effective for removing fMRI gradient and/or brain stimulation artifacts originating outside the head. Although beyond the scope of this paper, our preliminary tests on concurrent fMRI-EEG and EEG-NIBS recordings indicated successful removal of very high-amplitude electromagnetic noise components. Therefore, GEDAI’s automated, robust denoising capabilities based on biophysical principles could make it a promising candidate for improving data quality in challenging multimodal and stimulation experiments. Finally, as GEDAI utilizes a shared M/EEG forward model, its potential applicability could also extend to magnetoencephalography (MEG) recordings, especially optically-pumped magnetometers (OPM-MEG) where sensors maintain a fixed position relative to the head (Brookes et al., 2022).

### Limitations

First, GEDAI’s performance is highly dependent on an accurate match between the theoretical leadfield model and the actual EEG electrode positions. Any spatial mismatch will proportionally degrade its efficacy, making precise electrode placement a prerequisite for optimal results.

Second, because GEDAI defines artifacts as signals originating outside the brain, it is not designed to remove artifacts that originate from within the brain volume, such as those from deep brain stimulation, for example.

Third, the current implementation assumes uniform and uncorrelated brain sources. Incorporating a more neurophysiologically realistic source covariance model (e.g. one that accounts for functional networks and differing regional power) could improve performance, though this requires further research.

Finally, like PCA/ICA, the number of removable artifact components is limited by the number of channels, making GEDAI unsuitable for single-channel EEG.

## Conclusion

Taken together, these findings position GEDAI as a state-of-the-art tool for EEG denoising, distinguished by its robust performance in heavily contaminated, low-SNR conditions and its particular strength in resolving complex mixtures of artifacts.

## Data and code availability

The open-source GEDAI plugin for EEGLAB is available at: https://github.com/neurotuning/GEDAI-master

All simulated datasets and analysis codes used in this paper will be released as a standalone resource upon publication, known as “**BEAR: Benchmarking EEG Artifact Removal with Synthetic and Empirical Datasets**”. The open EEGLAB code will allow users to consult and modify the specific default parameters used in this study.

## Acknowledgements

We would like to thank Michael X Cohen for his helpful discussions as well as tutorials on EEG and GEVD, which have inspired this work. This study was supported by the Swiss National Science Foundation (SNSF), grant number 215712.

## Supplementary Results

**Fig S1:**
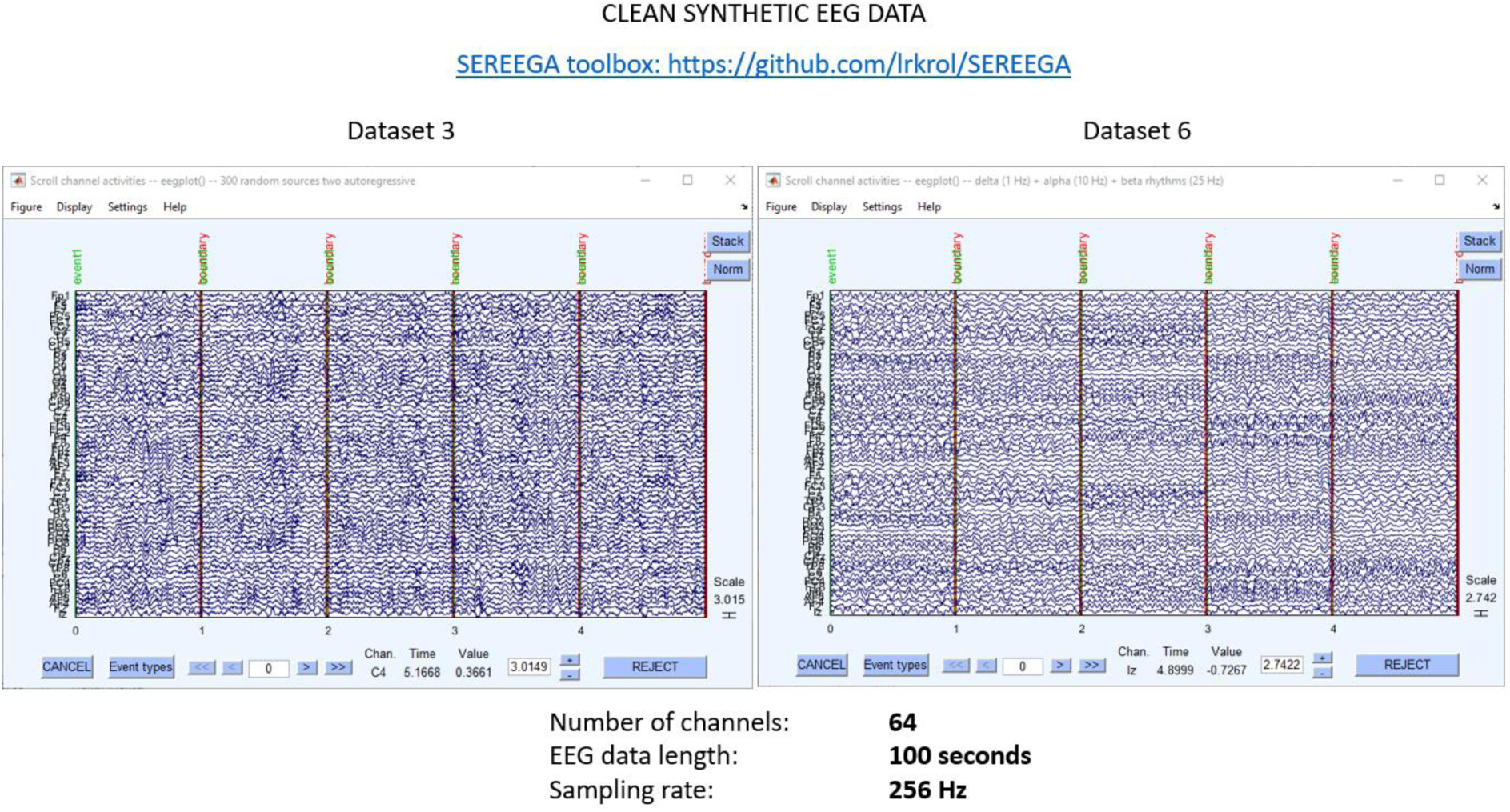
Clean *synthetic* data: example simulated ‘background’ EEG (i.e.noise-free) simulated with https://github.com/lrkrol/SEREEGA

**Fig S2.**
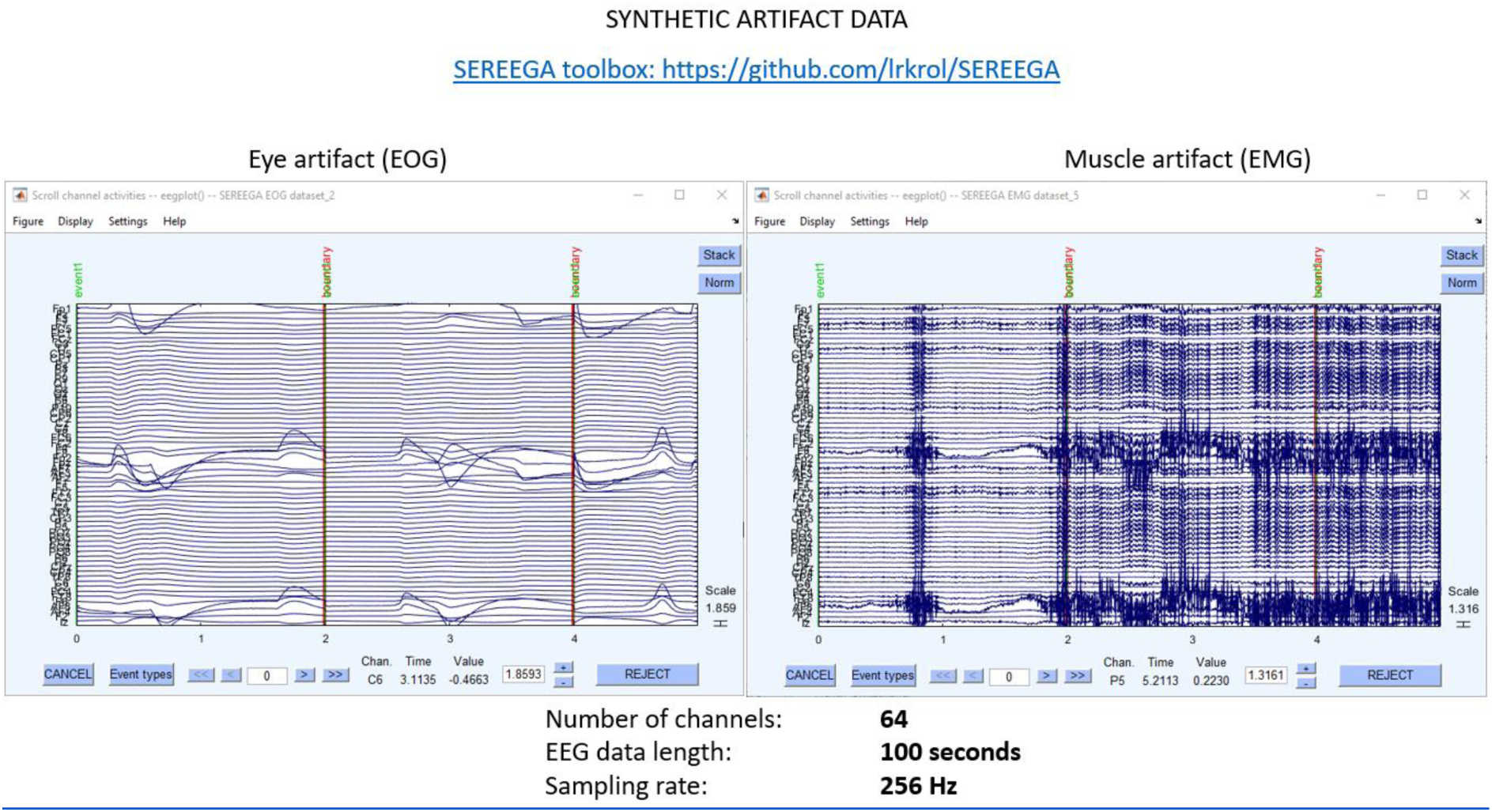
Artifactual *synthetic* data. : example of simulated data containing only EOG (left panel) or EMG artifacts (right panel) - using source locations from Fig S5 with https://github.com/lrkrol/SEREEGA + https://github.com/ncclabsustech/EEGdenoiseNet

**Fig S3.**
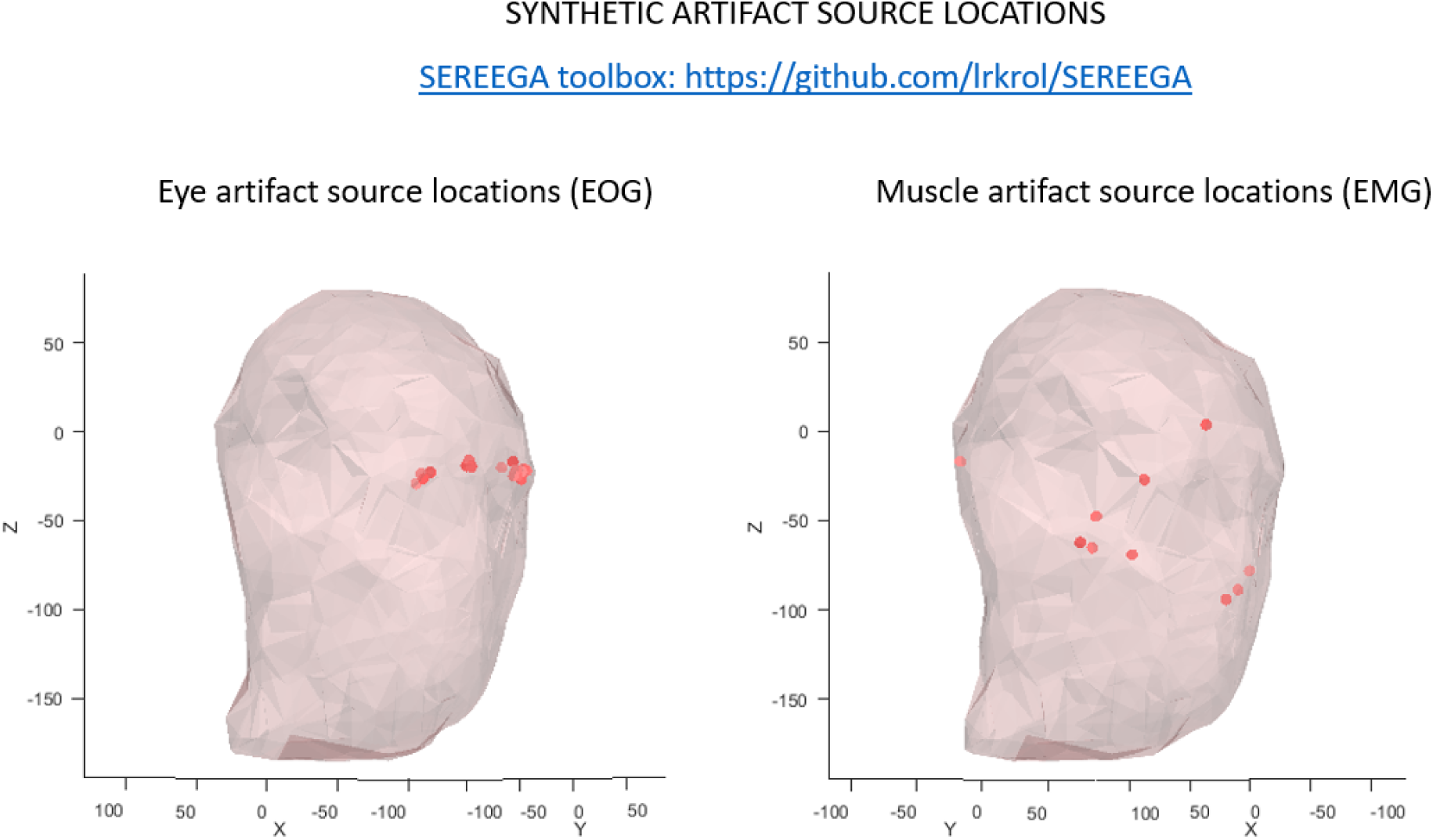
*Synthetic* artifact source locations for simulated EOG (left panel) or EMG artifacts (right panel) from https://github.com/lrkrol/SEREEGA

**Fig S4:**
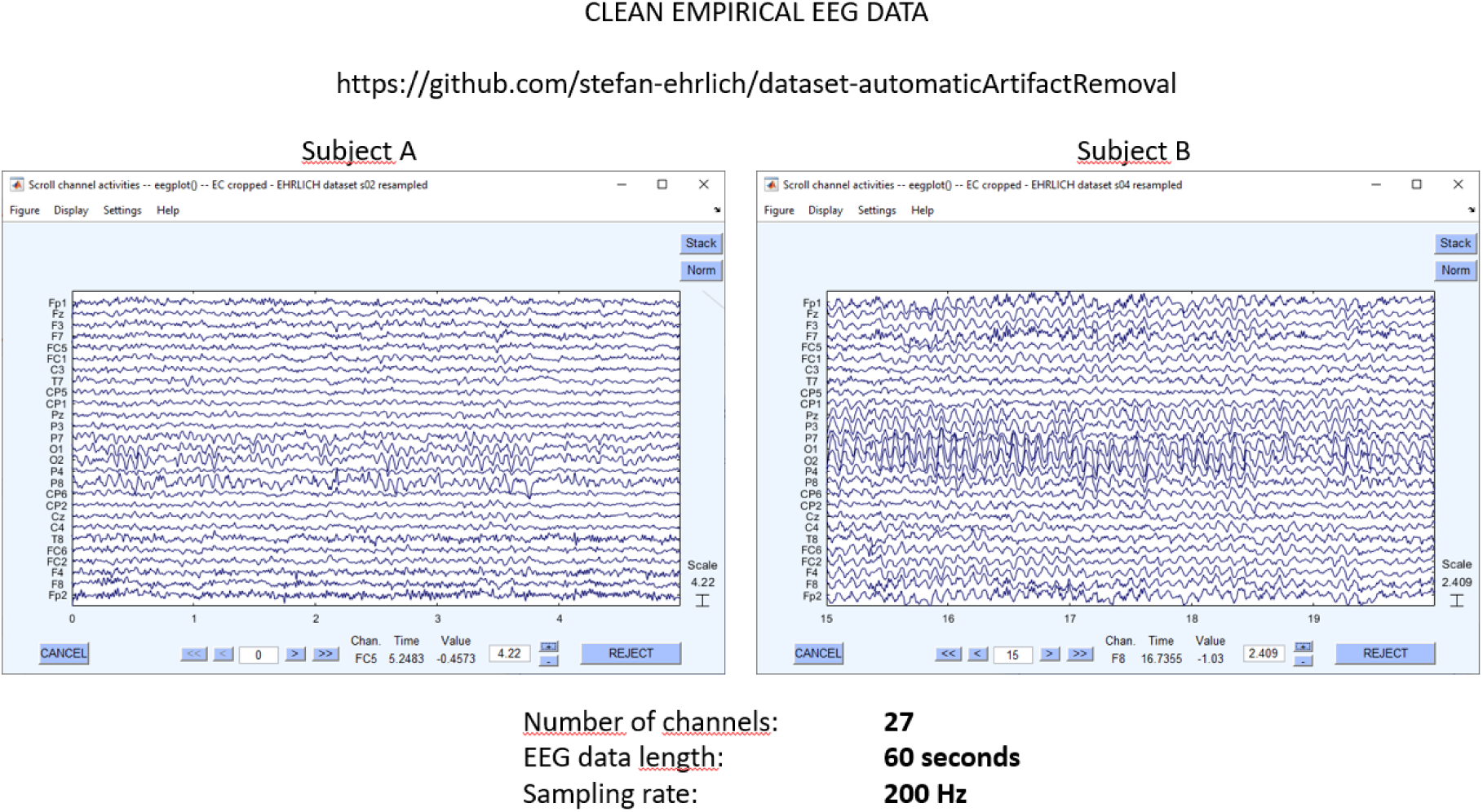
Clean empirical data. : example resting-state EEG recordings (i.e.noise-free) from https://github.com/stefan-ehrlich/dataset-automaticArtifactRemoval

**Fig S5.**
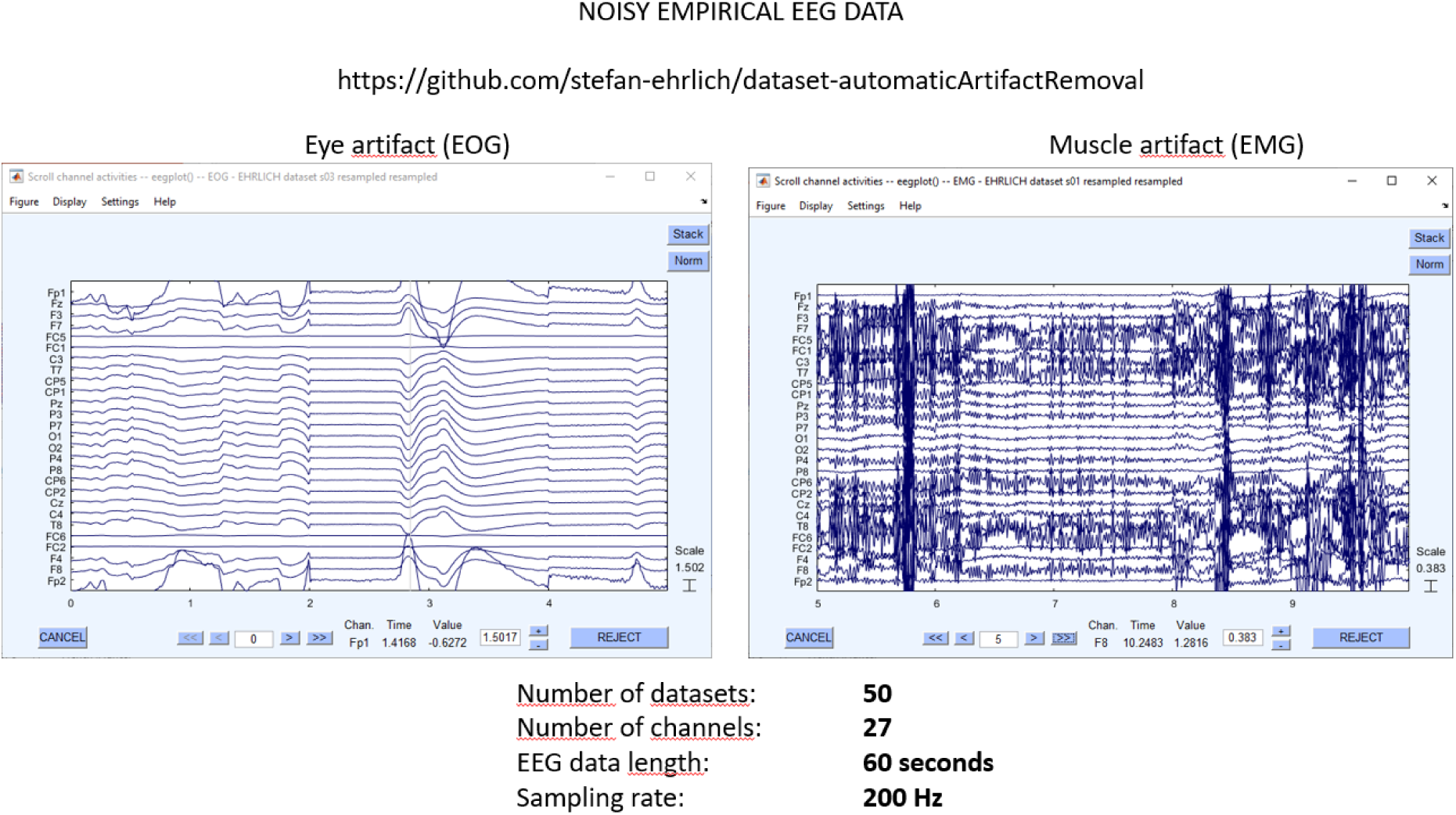
Artifactual *empirical* EEG data. : example of simulated data containing only EOG (left panel) or EMG artifacts (right panel) from https://github.com/stefan-ehrlich/dataset-automaticArtifactRemoval + https://github.com/ncclabsustech/EEGdenoiseNet

**Fig S6.**
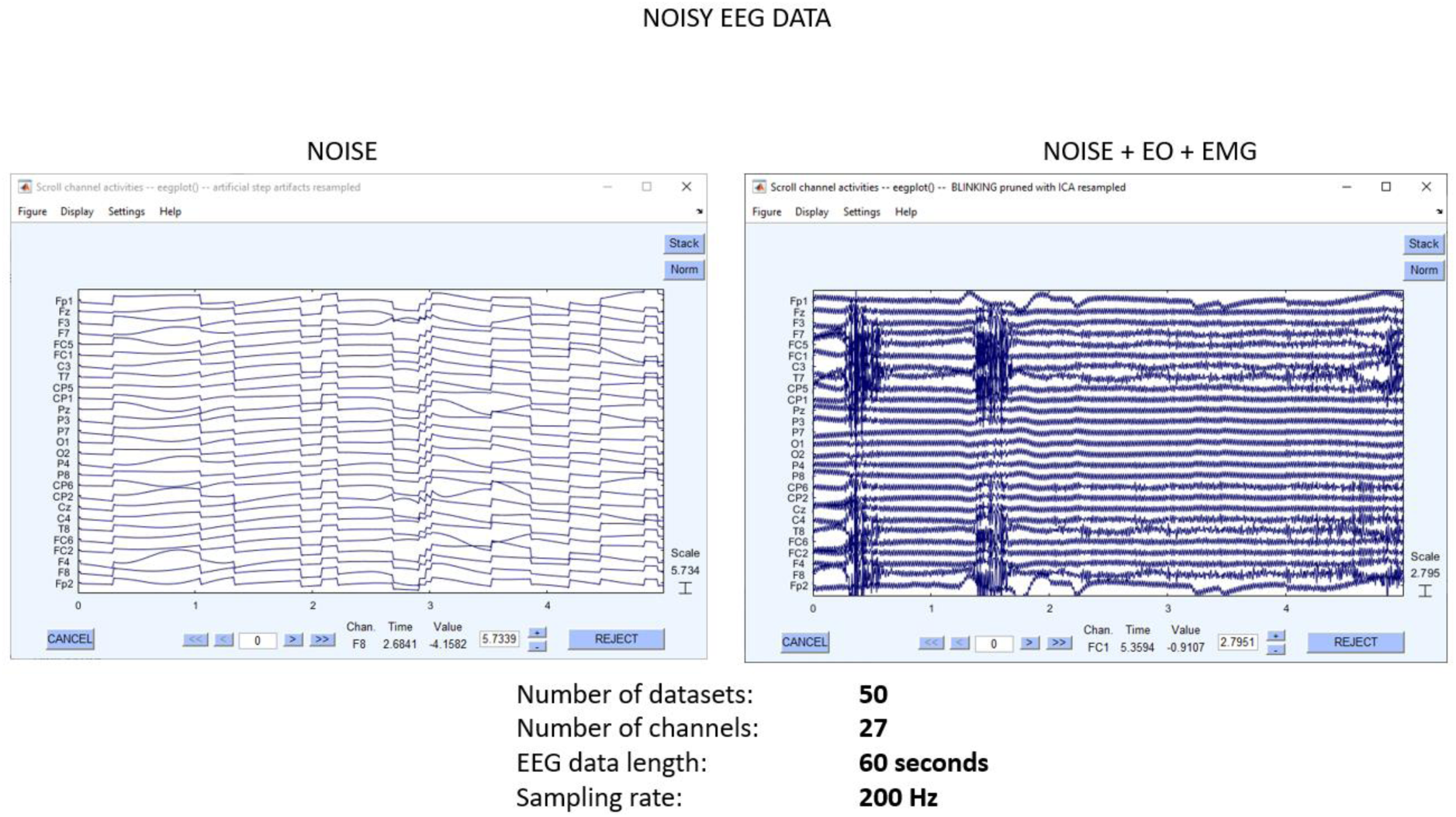
Noisy *empirical* EEG data. : example of simulated data containing step artifacts (left panel) or line noise from https://github.com/stefan-ehrlich/dataset-automaticArtifactRemoval + https://github.com/ncclabsustech/EEGdenoiseNet https://github.com/vpKumaravel/NEAR/tree/main/SimulateArtifacts

**Figure S7.**
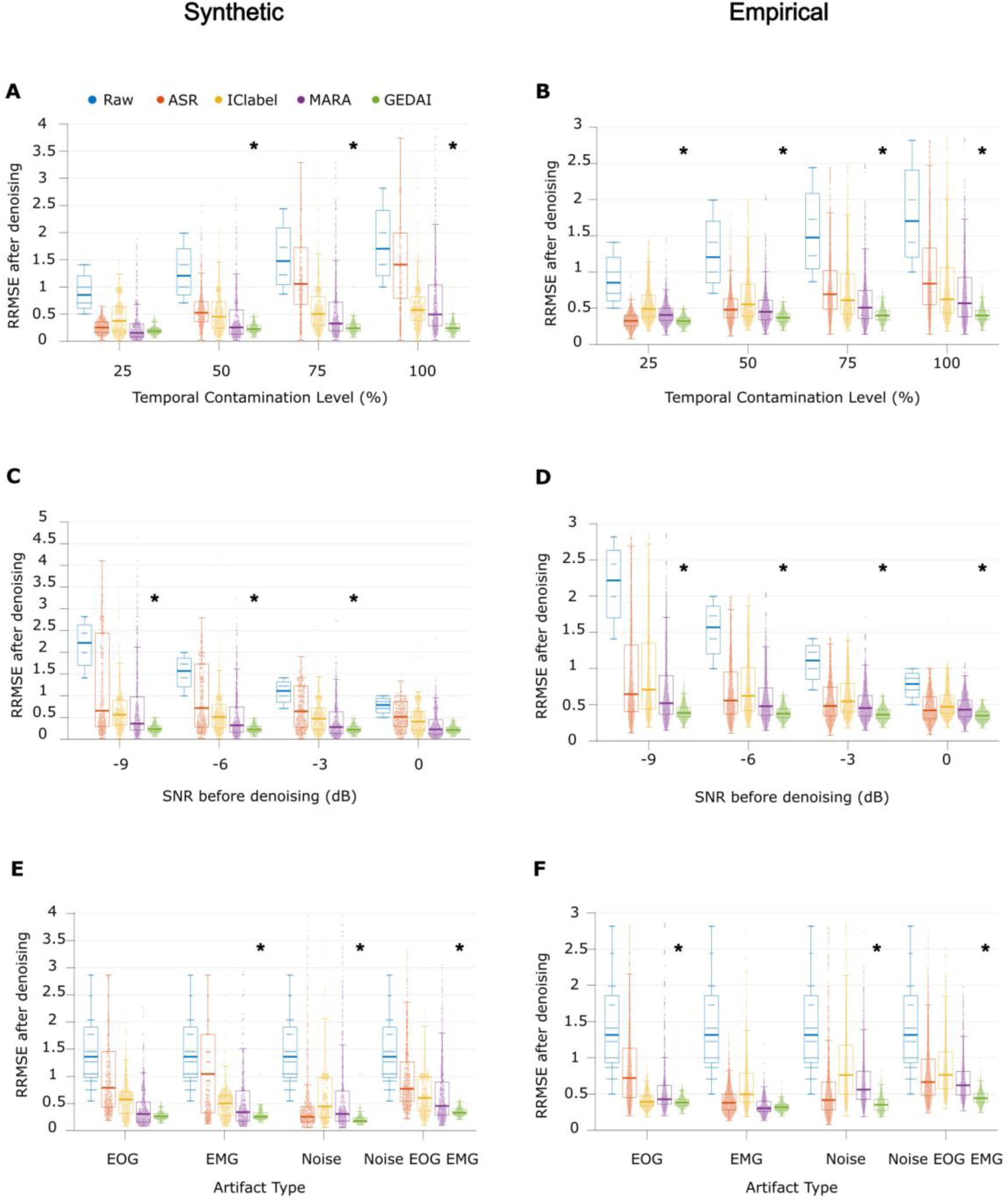
**Relative Root Mean Square Error (RRMSE)**. Lower values indicate better denoising. Asterisks indicate the winning algorithm in each case. No asterisk signifies a statistical tie. **Synthetic denoising by temporal contamination.** For every temporal contamination level, each coloured data point represents 1 of 800 datasets (50 datasets x 4 SNR levels x 4 artifact types). **Empirical denoising by temporal contamination.** Results are pooled across all baseline SNRbefore a*nd* all artifact types. For every temporal contamination level, each coloured data point represents 1 of 1,600 datasets (100 datasets x 4 SNR levels x 4 artifact types). **Synthetic denoising by SNRbefore.** Results are pooled across all temporal contamination levels and all artifact types. For every SNRbefore, each coloured data point represents 1 of 800 datasets (50 datasets x 4 artifact types x 4 temporal contamination levels). **Empirical denoising by SNRbefore**. Results are pooled across all temporal contamination levels and all artifact types. For every SNRbefore, each coloured data point represents 1 of 1,600 datasets (100 datasets x 4 artifact types x 4 temporal contamination levels). **Synthetic denoising by artifact type.** Results are pooled across all SNRbefore and temporal contamination levels. For every artifact type, each coloured data point represents 1 of 800 datasets (50 datasets x 4 SNR levels x 4 temporal contamination levels). **Empirical denoising by artifact type.** Results are pooled across all SNRbefore and temporal contamination levels. For every artifact type, each coloured data point represents 1 of 1,600 datasets (100 datasets x 4 SNR levels x 4 temporal contamination levels).

## Bibliography

1. Amico, E., & Goñi, J. (2018). The quest for identifiability in human functional connectomes. Scientific Reports, 8(1), 8254. 10.1038/s41598-018-25089-1

2. Amin, U., Nascimento, F. A., Karakis, I., Schomer, D., & Benbadis, S. R. (2023). Normal variants and artifacts : Importance in EEG interpretation. Epileptic Disorders, 25(5), 591- 648. 10.1002/epd2.20040

3. Bailey, N. W., Biabani, M., Hill, A. T., Miljevic, A., Rogasch, N. C., McQueen, B., Murphy, O. W., & Fitzgerald, P. B. (2023). Introducing RELAX : An automated pre-processing pipeline for cleaning EEG data - Part 1: Algorithm and application to oscillations. Clinical Neurophysiology, 149, 178- 201. 10.1016/j.clinph.2023.01.017

4. Bigdely-Shamlo, N., Mullen, T., Kothe, C., Su, K.-M., & Robbins, K. A. (2015). The PREP pipeline : Standardized preprocessing for large-scale EEG analysis. Frontiers in Neuroinformatics, 9, 16. 10.3389/fninf.2015.00016

5. Blankertz, B., Tomioka, R., Lemm, S., Kawanabe, M., & Muller, K. (2008). Optimizing Spatial filters for Robust EEG Single-Trial Analysis. IEEE Signal Processing Magazine, 25(1), 41- 56. 10.1109/MSP.2008.4408441

6. Boudet, S., Peyrodie, L., Forzy, G., Pinti, A., Toumi, H., & Gallois, P. (2012). Improvements of Adaptive Filtering by Optimal Projection to filter different artifact types on long duration EEG recordings. Computer Methods and Programs in Biomedicine, 108(1), 234- 249. 10.1016/j.cmpb.2012.04.005

7. Brookes, M. J., Leggett, J., Rea, M., Hill, R. M., Holmes, N., Boto, E., & Bowtell, R. (2022). Magnetoencephalography with optically pumped magnetometers (OPM-MEG) : The next generation of functional neuroimaging. Trends in Neurosciences, 45(8), 621- 634. 10.1016/j.tins.2022.05.008

8. Chang, C.-Y., Hsu, S.-H., Pion-Tonachini, L., & Jung, T.-P. (2020). Evaluation of Artifact Subspace Reconstruction for Automatic Artifact Components Removal in Multi- Channel EEG Recordings. IEEE Transactions on Biomedical Engineering, 67(4), 1114- 1121. 10.1109/TBME.2019.2930186

9. Clark, I., Biscay, R., Echeverría, M., & Virués, T. (1995). Multiresolution decomposition of non-stationary EEG signals : A preliminary study. Computers in Biology and Medicine, 25(4), 373- 382. 10.1016/0010-4825(95)00014-u

10. Cohen, M. X. (2022a). A tutorial on generalized eigendecomposition for denoising, contrast enhancement, and dimension reduction in multichannel electrophysiology. NeuroImage, 247, 118809. 10.1016/j.neuroimage.2021.118809

11. Cohen, M. X. (2022b). A tutorial on generalized eigendecomposition for denoising, contrast enhancement, and dimension reduction in multichannel electrophysiology. NeuroImage, 247, 118809. 10.1016/j.neuroimage.2021.118809

12. de Cheveigné, A., & Parra, L. C. (2014). Joint decorrelation, a versatile tool for multichannel data analysis. NeuroImage, 98, 487- 505. 10.1016/j.neuroimage.2014.05.068

13. DeLong, E. R., DeLong, D. M., & Clarke-Pearson, D. L. (1988). Comparing the areas under two or more correlated receiver operating characteristic curves : A nonparametric approach. Biometrics, 44(3), 837- 845.

14. Delorme, A., & Makeig, S. (2004). EEGLAB : An open source toolbox for analysis of single-trial EEG dynamics including independent component analysis. Journal of Neuroscience Methods, 134(1), 9- 21. 10.1016/j.jneumeth.2003.10.009

15. Delorme, A., Palmer, J., Onton, J., Oostenveld, R., & Makeig, S. (2012). Independent EEG sources are dipolar. PLoS ONE, 7(2), e30135. 10.1371/journal.pone.0030135

16. de Munck, J. C., van Dijk, B. W., & Spekreijse, H. (1988). Mathematical dipoles are adequate to describe realistic generators of human brain activity. IEEE Transactions on Biomedical Engineering, 35(11), 960- 966. 10.1109/10.8677

17. de Munck, J. C., Vijn, P. C., & da Silva, F. L. (1992). A random dipole model for spontaneous brain activity. IEEE transactions on biomedical engineering, 39(8), 791- 804.

18. Ehrlich, S. (2024). *Stefan-ehrlich/dataset-automaticArtifactRemoval* [Logiciel]. https://github.com/stefan-ehrlich/dataset-automaticArtifactRemoval (Édition originale 2019)

19. Fratangelo, R., Lolli, F., Scarpino, M., & Grippo, A. (2025). Point-of-Care Electroencephalography in Acute Neurological Care : A Narrative Review. Neurology International, 17(4), Article 4. 10.3390/neurolint17040048

20. Frølich, L., Andersen, T. S., & Mørup, M. (2015). Classification of independent components of EEG into multiple artifact classes. Psychophysiology, 52(1), 32- 45. 10.1111/psyp.12290

21. Gabard-Durnam, L. J., Mendez Leal, A. S., Wilkinson, C. L., & Levin, A. R. (2018). The Harvard Automated Processing Pipeline for Electroencephalography (HAPPE) : Standardized Processing Software for Developmental and High-Artifact Data. Frontiers in Neuroscience, 12. 10.3389/fnins.2018.00097

22. Gouy-Pailler, C., Sameni, R., Congedo, M., & Jutten, C. (2009). Iterative Subspace Decomposition for Ocular Artifact Removal from EEG Recordings. In T. Adali, C. Jutten, J. M. T. Romano, & A. K. Barros (Éds.), Independent Component Analysis and Signal Separation (Vol. 5441, p. 419- 426). Springer Berlin Heidelberg. 10.1007/978-3-642-00599-2_53

23. Graichen, U., Eichardt, R., Fiedler, P., Strohmeier, D., Zanow, F., & Haueisen, J. (2015). SPHARA - A Generalized Spatial Fourier Analysis for Multi-Sensor Systems with Non-Uniformly Arranged Sensors : Application to EEG. PLOS ONE, 10(4), e0121741. 10.1371/journal.pone.0121741

24. Gramfort, A., Papadopoulo, T., Olivi, E., & Clerc, M. (2010). OpenMEEG: opensource software for quasistatic bioelectromagnetics. BioMedical Engineering OnLine, 9(1), 45. 10.1186/1475-925X-9-45

25. Hajhassani, D., Mattout, J., & Congedo, M. (2024). An Automatic Riemannian Artifact Rejection Method for P300-based BCIs. 2024 *32nd European Signal Processing Conference (EUSIPCO)*, 1616- 1620. 10.23919/EUSIPCO63174.2024.10715349

26. Harmening, N., Klug, M., Gramann, K., & Miklody, D. (2022). HArtMuT—modeling eye and muscle contributors in neuroelectric imaging. Journal of Neural Engineering, 19(6), 066041. 10.1088/1741-2552/aca8ce

27. Hartmann, M. M., Schindler, K., Gebbink, T. A., Gritsch, G., & Kluge, T. (2014). PureEEG: Automatic EEG artifact removal for epilepsy monitoring. Neurophysiologie clinique = Clinical neurophysiology, 44(5), 479- 490. 10.1016/j.neucli.2014.09.001

28. Haslacher, D., Nasr, K., Robinson, S. E., Braun, C., & Soekadar, S. R. (2021). Stimulation artifact source separation (SASS) for assessing electric brain oscillations during transcranial alternating current stimulation (tACS). NeuroImage, 228, 117571. 10.1016/j.neuroimage.2020.117571

29. Hsu, S.-H., Mullen, T., Jung, T.-P., & Cauwenberghs, G. (2014). Online recursive independent component analysis for real-time source separation of high-density EEG 2014 36t h Annual International Conference of the IEEE Engineering in Medicine and Biology Society, 3845- 3848. 10.1109/EMBC.2014.6944462

30. Huang, Y., Parra, L. C., & Haufe, S. (2016). The New York Head-A precise standardized volume conductor model for EEG source localization and tES targeting. NeuroImage, 140, 150- 162. 10.1016/j.neuroimage.2015.12.019

31. Ille, N., Berg, P., & Scherg, M. (2002). Artifact correction of the ongoing EEG using spatial filters based on artifact and brain signal topographies. Journal of Clinical Neurophysiology: Official Publication of the American Electroencephalographic Society, 19(2), 113- 124. 10.1097/00004691-200203000-00002

32. Jung, T. P., Makeig, S., Humphries, C., Lee, T. W., McKeown, M. J., Iragui, V., & Sejnowski, T. J. (2000). Removing electroencephalographic artifacts by blind source separation. Psychophysiology, 37(2), 163- 178.

33. Knyazev, A. V., & Argentati, M. E. (2002). Principal Angles between Subspaces in an A- Based Scalar Product : Algorithms and Perturbation Estimates. SIAM Journal on Scientific Computing, 23(6), 2008- 2040. 10.1137/S1064827500377332

34. Koles, Z. J. (1991). The quantitative extraction and topographic mapping of the abnormal components in the clinical EEG. Electroencephalography and Clinical Neurophysiology, 79(6), 440- 447. 10.1016/0013-4694(91)90163-x

35. Kothe, C. A. E. (2013). The artifact subspace reconstruction method. Accessed: Jul, 17, 2017.

36. Kothe, C. A. E., & Jung, T.-P. (2015). *Artifact removal techniques with signal reconstruction* (World Intellectual Property Organization Brevet No. WO2015047462A9). https://patents.google.com/patent/WO2015047462A9/en

37. Krol, L. R., Pawlitzki, J., Lotte, F., Gramann, K., & Zander, T. O. (2018). SEREEGA : Simulating event-related EEG activity. Journal of Neuroscience Methods, 309, 13- 24. 10.1016/j.jneumeth.2018.08.001

38. Kumaravel, V. P., Farella, E., Parise, E., & Buiatti, M. (2022). NEAR : An artifact removal pipeline for human newborn EEG data. Developmental Cognitive Neuroscience, 54, 101068. 10.1016/j.dcn.2022.101068

39. Lazarevic, A., & Kumar, V. (2005). Feature bagging for outlier detection. Proceedings of the eleventh ACM SIGKDD international conference on Knowledge discovery in data mining, 157- 166. 10.1145/1081870.1081891

40. Lee, T.-W., Girolami, M., & Sejnowski, T. J. (1999). Independent Component Analysis Using an Extended Infomax Algorithm for Mixed Subgaussian and Supergaussian Sources. Neural Computation, 11(2), 417- 441. 10.1162/089976699300016719

41. Li, Z., Zhao, Y., Hu, X., Botta, N., Ionescu, C., & Chen, G. H. (2023). ECOD : Unsupervised Outlier Detection Using Empirical Cumulative Distribution Functions. IEEE Transactions on Knowledge and Data Engineering, 35(12), 12181- 12193. 10.1109/TKDE.2022.3159580

42. Lotte, F., & Guan, C. (2011). Regularizing common spatial patterns to improve BCI designs : Unified theory and new algorithms. IEEE Transactions on Bio-Medical Engineering, 58(2), 355- 362. 10.1109/TBME.2010.2082539

43. Mak, J. N., & Wolpaw, J. R. (2009). Clinical Applications of Brain-Computer Interfaces : Current State and Future Prospects. IEEE reviews in biomedical engineering, 2, 187- 199. 10.1109/RBME.2009.2035356

44. Mumtaz, W., Rasheed, S., & Irfan, A. (2021). Review of challenges associated with the EEG artifact removal methods. Biomedical Signal Processing and Control, 68, 102741. 10.1016/j.bspc.2021.102741

45. Mushtaq, F., Welke, D., Gallagher, A., Pavlov, Y. G., Kouara, L., Bosch-Bayard, J., van den Bosch, J. J. F., Arvaneh, M., Bland, A. R., Chaumon, M., Borck, C., He, X., Luck, S. J., Machizawa, M. G., Pernet, C., Puce, A., Segalowitz, S. J., Rogers, C., Awais, M., … Valdes-Sosa, P. (2024). One hundred years of EEG for brain and behaviour research. Nature Human Behaviour, 8(8), 1437- 1443. 10.1038/s41562-024-01941-5

46. Mutanen, T. P., Metsomaa, J., Liljander, S., & Ilmoniemi, R. J. (2018). Automatic and robust noise suppression in EEG and MEG : The SOUND algorithm. NeuroImage, 166, 135- 151. 10.1016/j.neuroimage.2017.10.021

47. Olbrich, S., Jödicke, J., Sander, C., Himmerich, H., & Hegerl, U. (2011). ICA-based muscle artefact correction of EEG data : What is muscle and what is brain?: Comment on McMenamin et al. NeuroImage, 54(1), 1- 3. 10.1016/j.neuroimage.2010.04.256

48. Omejc, N., Peskar, M., Miladinović, A., Kavcic, V., Džeroski, S., & Marusic, U. (2023). On the Influence of Aging on Classification Performance in the Visual EEG Oddball Paradigm Using Statistical and Temporal Features. Life, 13(2), Article 2. 10.3390/life13020391

49. Parra, L., & Sajda, P. (2003). Blind Source Separation via Generalized Eigenvalue Decomposition. Journal of Machine Learning Research, 4, 1261- 1269.

50. Pion-Tonachini, L., Kreutz-Delgado, K., & Makeig, S. (2019). ICLabel : An automated electroencephalographic independent component classifier, dataset, and website. NeuroImage, 198, 181- 197. 10.1016/j.neuroimage.2019.05.026

51. Radüntz, T., Scouten, J., Hochmuth, O., & Meffert, B. (2017). Automated EEG artifact elimination by applying machine learning algorithms to ICA-based features. Journal of Neural Engineering, 14(4), 046004. 10.1088/1741-2552/aa69d1

52. Ros, T., Baars, B. J., Lanius, R. A., & Vuilleumier, P. (2014). Tuning pathological brain oscillations with neurofeedback : A systems neuroscience framework. Frontiers in Human Neuroscience, 8, 1008. 10.3389/fnhum.2014.01008

53. Sawangjai, P., Hompoonsup, S., Leelaarporn, P., Kongwudhikunakorn, S., & Wilaiprasitporn, T. (2020). Consumer Grade EEG Measuring Sensors as Research Tools : A Review.

54. *IEEE Sensors Journal*, *20*(8), 3996- 4024. IEEE Sensors Journal. 10.1109/JSEN.2019.2962874

55. Somers, B., Francart, T., & Bertrand, A. (2018). A generic EEG artifact removal algorithm based on the multi-channel Wiener filter. Journal of Neural Engineering, 15(3), 036007. 10.1088/1741-2552/aaac92

56. Sorrentino, P., Rucco, R., Lardone, A., Liparoti, M., Troisi Lopez, E., Cavaliere, C., Soricelli, A., Jirsa, V., Sorrentino, G., & Amico, E. (2021). Clinical connectome fingerprints of cognitive decline. NeuroImage, 238, 118253. 10.1016/j.neuroimage.2021.118253

57. Tadel, F., Baillet, S., Mosher, J. C., Pantazis, D., & Leahy, R. M. (2011). Brainstorm : A User-Friendly Application for MEG/EEG Analysis. Computational Intelligence and Neuroscience, 2011, 879716. 10.1155/2011/879716

58. Taulu, S., Kajola, M., & Simola, J. (2004). Suppression of interference and artifacts by the Signal Space Separation Method. Brain Topography, 16(4), 269- 275. 10.1023/b:brat.0000032864.93890.f9

59. Uusitalo, M. A., & Ilmoniemi, R. J. (1997). Signal-space projection method for separating MEG or EEG into components. Medical & Biological Engineering & Computing, 35(2), 135- 140. 10.1007/BF02534144

60. Wang, F., Ma, Y., Gao, T., Tao, Y., Wang, R., Zhao, R., Cao, F., Gao, Y., & Ning, X. (2025). Repairbads : An automatic and adaptive method to repair bad channels and segments for OPM-MEG. NeuroImage, 306, 120996. 10.1016/j.neuroimage.2024.120996

61. Wang, Y., Berg, P., & Scherg, M. (1999). Common spatial subspace decomposition applied to analysis of brain responses under multiple task conditions : A simulation study. Clinical Neurophysiology: Official Journal of the International Federation of Clinical Neurophysiology, 110(4), 604- 614. 10.1016/s1388-2457(98)00056-x

62. Weinstein, D., Zhukov, L., & Johnson, C. (2000). Lead-field bases for electroencephalography source imaging. Annals of Biomedical Engineering, 28(9), 1059- 1065. 10.1114/1.1310220

63. Winkler, I., Brandl, S., Horn, F., Waldburger, E., Allefeld, C., & Tangermann, M. (2014). Robust artifactual independent component classification for BCI practitioners. Journal of Neural Engineering, 11(3), 035013. 10.1088/1741-2560/11/3/035013

64. Zhang, H., Zhao, M., Wei, C., Mantini, D., Li, Z., & Liu, Q. (2021). EEGdenoiseNet : A benchmark dataset for deep learning solutions of EEG denoising. Journal of Neural Engineering, 18(5), 056057. 10.1088/1741-2552/ac2bf8

65. Zhang, X. (2024). Deep Learning-Based Techniques for Electroencephalogram (EEG) Signal Denoising. Transactions on Computer Science and Intelligent Systems Research, 5, 922- 927. 10.62051/rrve8560

